# Cortical reliability amid noise and chaos

**DOI:** 10.1101/304121

**Authors:** Max Nolte, Michael W. Reimann, James G. King, Henry Markram, Eilif B. Muller

## Abstract

Typical responses of cortical neurons to identical sensory stimuli are highly variable. It has thus been proposed that the cortex primarily uses a rate code. However, other studies have argued for spike-time coding under certain conditions. The potential role of spike-time coding is constrained by the intrinsic variability of cortical circuits, which remains largely unexplored. Here, we quantified this intrinsic variability using a biophysical model of rat neocortical microcircuitry with biologically realistic noise sources. We found that stochastic neurotransmitter release is a critical component of this variability, which, amplified by recurrent connectivity, causes rapid chaotic divergence with a time constant on the order of 10-20 milliseconds. Surprisingly, weak thalamocortical stimuli can transiently overcome the chaos, and induce reliable spike times with millisecond precision. We show that this effect relies on recurrent cortical connectivity, and is not a simple effect of feed-forward thalamocortical input. We conclude that recurrent cortical architecture supports millisecond spike-time reliability amid noise and chaotic network dynamics, resolving a long-standing debate.

## Introduction

The typical electrical activity of cortical neurons is highly variable, in the sense that membrane potentials, spike times and interspike intervals vary during spontaneous activity as well as across trials with identical sensory stimuli^1–4^. While part of this variability could be due to intrinsic noise sources, a substantial part could also be due to hidden variables such as unknown input from other parts of the brain, environmental parameters, or brain state^5,6^. For instance, it has been shown that, in the visual cortex, the act of running modulates responses of neurons to identical stimuli^7^. Moreover, some neurons in sensory cortices can encode sensory input with high spike-time precision^8–10^. Taken together, it is compelling to assume that intrinsic noise plays a negligible role, and that cortical variability is essentially deterministic^11^, encoding hidden or unobserved variables. This view is also supported by the fact that neocortical neurons respond to somatic current injections in vitro with high reliability^12^. However, there are two important reasons to believe that a large part of cortical variability is due to internally generated noise that carries no signal.

First, all cortical neurons are subject to well-established cellular noise sources, such as stochastic synaptic transmission and ion-channel noise^13^. These noise sources ultimately originate from proteins susceptible to thermodynamic fluctuations, and are therefore indeed truly intrinsic sources of noise^6,13^. In particular, synaptic transmission is based on a sequence of stochastic molecular events, where the low numbers of molecules involved do not allow stochastic properties to average out^14^. In fact, in tightly controlled slice environments in vitro, the probability of vesicle release upon action potential arrival at a single cortical synapse is low (∼50% between thick tufted layer 5 pyramidal neurons^15^), and estimated to be substantially lower in vivo^16^ (∼10% between same neurons^17^). The universal presence of synaptic noise suggests that cortical neurons respond far less reliably to presynaptic inputs than to current injections. It has been shown, moreover, that a simplified cortical network model with stochastic synapses can provide a sufficient explanation for variable spiking^18^. Furthermore, in vitro, some types of inhibitory neurons exhibit stochastic firing types. That is, they respond highly irregularly to constant somatic current injections^19^. This is due to ion-channel noise that is amplified during action potential initiation^20^. Even activity in regular firing excitatory neurons can be subject to ion-channel noise, for example during action potential propagation in thin axons^21^.

Second, models suggest^22,23^ and experiments show^24^ that cortical networks have chaotic dynamics. This implies, by definition, that small perturbations, such as those due to intrinsic cellular noise, are amplified. Thus, extra or missing spikes in the network, for example due to failed synaptic transmission, could fundamentally alter the trajectories of spiking activity in the network, leading in turn to large steady-state fluctuations.

In spite of their potential importance, the separate and combined impacts of network dynamics and cellular noise sources on cortical neuronal variability remain largely unexplored. There are several reasons why understanding what proportion of cortical neuronal variability is generated internally—and how this variability arises—is crucial for understanding the neural code.

First, strong internally generated variability due to chaotic network dynamics could prevent coding based on spike timing past the sensory periphery, and favor theories of firing rate coding^24^. To test the feasibility of models of cortical coding that rely on spike timing ^25–27^, we need to understand internal variability and how it arises.

Second, variability could carry information and encode signals itself, for example perceptual uncertainty ^28^. It is thus essential to understand how to separate intrinsically generated variability that is *bona fide* noise from variability that encodes an additional signal or brain state.

Third, and more generally, optimal coding strategies for neural circuits depend on where noise enters the circuit^29^. That is, to understand the neural code, we need to understand the mechanisms responsible for internally generated variability. Currently, it is impossible to measure all external inputs to a local population of cortical neurons in vivo. As a result, we are still unable to quantify how much of the experimentally observed variability is generated internally by the local circuitry, and how much is generated externally.

In this study, we addressed these questions with a recently developed simulation-based approach, namely a biologically constrained model of a prototypical *neocortical microcircuit* in rat somatosensory cortex (the NMC-model; see Markram *et al*.^17^). The NMC-model, which reproduces a range of in vivo experiments, incorporates several prominent sources of noise, including stochastic synaptic transmission and ion channel noise. Each of these noise sources is constrained to replicate experimentally observed variability. This bottom-up modeling approach provides control over all noise sources, as well as external inputs and internal states.

Through a series of simulation experiments, in which we selectively enabled noise sources and recurrent network dynamics, we characterized intrinsic cortical variability and how it arises. We confirmed that recurrent cortical dynamics are chaotic, but we found that an interplay of stochastic synaptic transmission and network dynamics determined the rate by which membrane potentials diverged. Surprisingly, the recurrent cortical circuitry can transiently overcome these chaotic network dynamics in response to weak thalamocortical inputs, and produce reliable spike timing. Thus, recurrent cortical architecture can transform relatively weak inputs into reliable patterns of activity amid high cellular noise and chaotic network dynamics.

## Results

### Rapid divergence of spontaneous activity

Using the NMC-model of rat somatosensory cortex (31’346 neurons, ∼8 million connections, and ∼36 million synapses; see **Figure 1a**), we simulated in vivo-like spontaneous neuronal activity. The NMC-model contains three types of biological noise sources, all of which are required to replicate neuronal responses to paired recordings and current injections in vitro (**Fig. 1b**). Each of the 36 million synapses in the model incorporates stochastic models of vesicle release, which display both *failure* of vesicle release (*a*) and *spontaneous release* (*b*). In this way, synaptic variability is biologically constrained. The irregular firing electrical neuron types (e-types) (1’137 neurons) also contain models of *stochastic potassium channels* (*c*), which induce irregular firing in response to constant current injections in vitro. A fourth, tunable noise source consisted of a noisy current (*d*) injected into the soma of each of the 31’346 neurons in the model, making it possible to depolarize neurons to in vivo-like levels (see Methods, Markram *et al*.^17^) (**Fig. 1b**). In our initial experiments, we maintained the magnitude of this generic noise far below the magnitude of the experimentally-constrained noise sources, using it later for sensitivity analysis. Realizations of the stochastic processes underlying these noise sources were determined by *sequences of random numbers*. By generating the sequences with different *random seeds*, we were able to obtain different, but equally valid probabilistic simulation trials.

**Figure 1.**
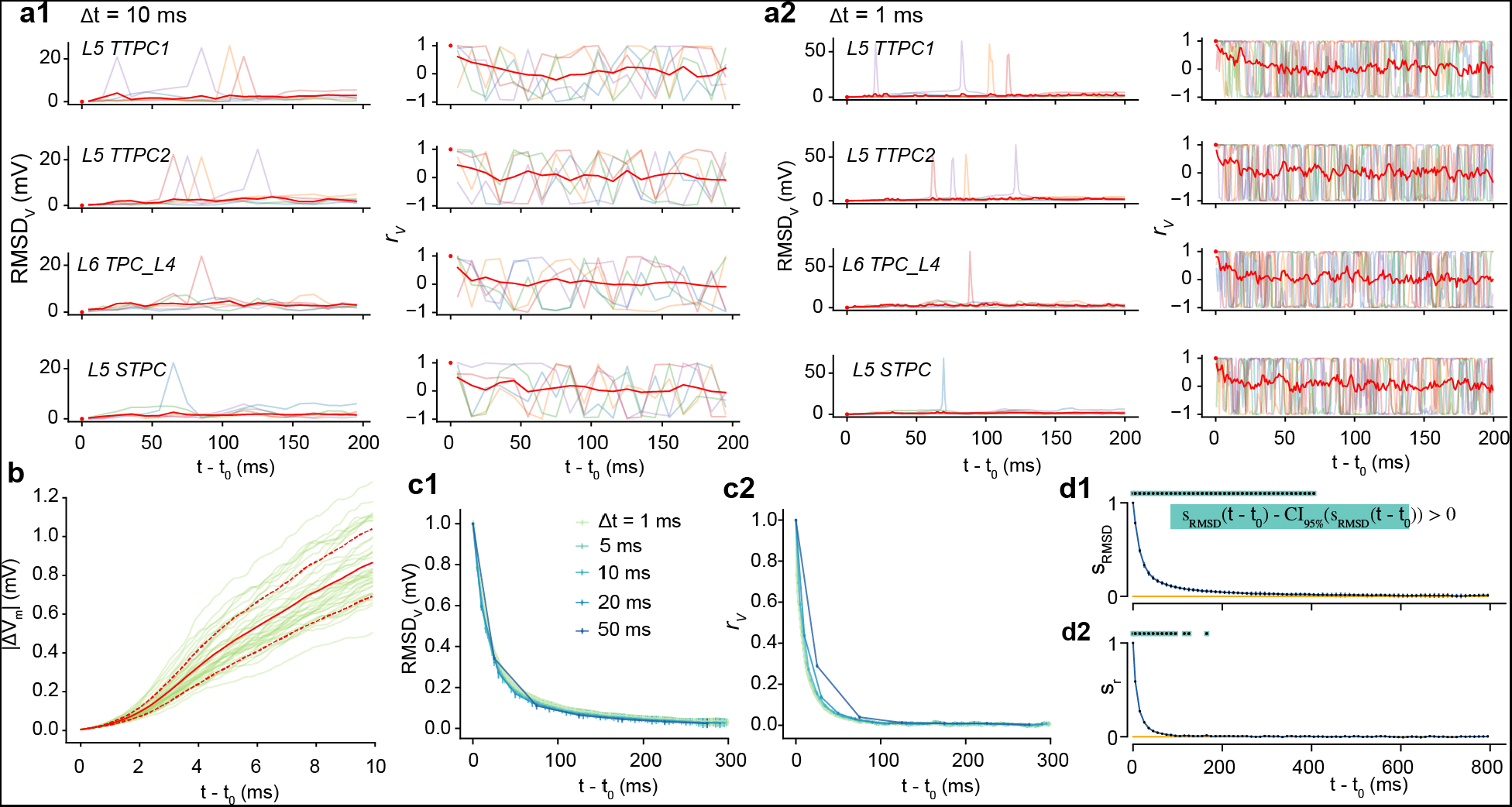
Rapid divergence of spontaneous activity. (**a**)Morphologically-detailed model of a neocortical microcircuit (NMC); depicted are 100 randomly selected neurons, out of 31’346 in total (∼0.3%). Neurons are colored according to their layer. (**b**) Examples of simulated noise sources in the NMC-model: stochastic synaptic transmission, including(*a*) vesicle release failure and (*b*) spontaneous vesicle release (‘miniature PSPs’) at all 36 million synapses; (*c*) probabilistic opening and closing of voltage-gated potassium channels in irregularly spiking inhibitory neurons (1’137 out of 31’346 neurons); (*d*) a constant depolarizing current with a weak white noise component 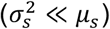 injected into the somata of all neurons. (**c**) The membrane potential of four sample neurons (and population mean of all 31’346 neurons) during a network simulation of in vivo-like spontaneous activity. At *t*_0_, the state of the microcircuit is saved, and then resumed twice with identical initial conditions, but with different random seeds for all noise sources. (**d**) Root-mean square deviation (*RMSD_v_*(*t*)) and correlation (*r_v_*(*t*)) of the somatic membrane potentials between pairs of resumed simulations diverging from identical initial conditions (mean of all neurons and 40 saved base states ± 95% confidence interval). The dashed lines depict the steady-state *RMSD_V_* and *r_v_* between independent simulations (i.e. resumed from different base states). (**e**) Time evolution of distributions of mean *RMSD_V_* and *r_v_* values for individual neurons. (**f**) The similarity of the system (*s_RMSD_* and *s_r_*) defined as the difference between the diverging and steady-state *RMSD_V_* and *r_v_*, normalized to lie between 1 (identical) and 0 (fully diverged) (mean ± 95% confidence interval). Exponential fit of *s_RM4D_* and *s_r_* for *t* - *t*_0_ < 40 ms (estimated time constant ± 68% confidence interval of fit).

Independent trials of electrical activity were simulated up to a time *t*_0_, at which point we saved the full dynamical state of the simulation (*base state*). We then *resumed* the simulation two times from the base state, i.e. we used identical initial conditions and histories in each case, but with different sequences of random numbers. This allowed us to obtain two equally valid probabilistic network trajectories for *t* > *t*_0_ for each base state. We observed that somatic membrane potentials (*V_m_*) for individual neurons, and the mean potentials for the population both diverged rapidly between the two simulations (**Fig. 1c**).

To quantify the time course of the divergence for each neuron *n*, we calculated the *root-mean-square deviation* of its somatic membrane potential in two trials in time bins of size ∆*t* starting from *t*_0_:

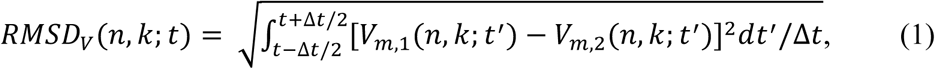

where *V_m,l_*(*n, k; t*) and *V*_*m*,2_(*n, k; t*) denote the time series of somatic membrane potentials of neuron n in two trials resuming from the same base state *k*. We consequently defined the mean root-mean-square deviation of the microcircuit *RMSD_v_*(*t*) as the mean of *RMSD_v_*(*n, k; t*) over all base states (*K=40*) and neurons (*N=31’346*). We observed that *RMSD_v_*(*t*) diverged rapidly from zero and eventually converged towards a steady-state value *RMSD*_∞_, equal to the *RMSD_v_* of independent trials that did not share the same base state (**Fig. 1d**, solid black and solid grey lines). The divergence was fast, with *RMSD_v_*(*t*) reaching more than 50% of its steady-state value within 20 ms.

While the *RMSD_v_*(*t*) of the circuit allowed us to accurately track the overall divergence of the whole circuit, *RMSD_v_*(*n, k; t*) for individual neurons and trials were too noisy for in-depth analysis (**Fig. 1e1** and **Supplementary Fig. 1a**). We note that while *RMSD_v_*(*t*) quantifies the absolute distance between membrane potentials, potentials can still be correlated independent of this distance. To this end, we also computed the *linear correlation* for each neuron for each base state, again for time bins of size ∆*t*:

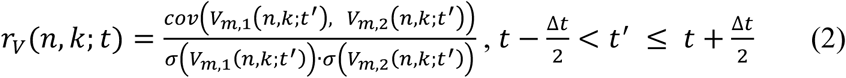

We found that the mean correlation *r_v_*(*t*) diverged faster than the absolute distance as measured by *RMSD_v_*(*t*) (**Fig. 1d**, dashed blue line), again with a broad distribution across individual neurons (**Fig. 1e2** and **Supplementary Fig. 1a**).

To better evaluate the difference between *r_v_*(*t*) and *RMSD_v_*(*t*), we computed the *similarity s_RMSD_*(*t*) of the microcircuit activity as the normalized difference between diverging and steady-state *RMSD_v_*(*t*) (and similarly *s_r_*(*t*) for *r_v_*(*t*)). When similarity s_*RMSD*_(*t*) = 1, membrane potential traces are identical; when *s_RMSD_*(*t*) =0 membrane potentials have reached their steady-state distance *RMSD*_∞_. Similarly, when *s_r_*(*t*) =1, membrane potentials have a perfect linear relationship; when *s_r_*(*t*) =0, they reached their steady-state correlation *r*_∞_. Comparing *s_r_*(*t*) and *s_RMSD_*(*t*), we observed that *r_v_*(*t*) diverged approximately twice as fast as *RMSD_v_*(*t*) (**Fig. 1f1** vs. **Fig. 1f2**). More precisely, an exponential fit to the first 40 milliseconds revealed divergence time constants of *τ _RMSD_* = 22.7 + 0.5 ms and *τ_r_* = 11.5 + 0.2 ms (± 68% confidence interval of fit). These were conserved for different bins sizes ∆*t*, with similar values for bin sizes ranging from 1 ms to 50 ms (**Supplementary Fig. 1c1,2**). We observe, however, that simple exponential decay does not provide an adequate description of the whole time-course of the similarity, as the time constant changes continuously, especially in the first several milliseconds (**Supplementary Fig. 1b**). While the initial divergence is rapid, a small, but statistically significant difference (p < 0.025) between diverging and independent activity persists for around 400 ms for *RMSD_v_* (**Supplementary Fig. 1d1**) and around 200 ms for *r_v_* (**Supplementary Fig. 1d2**).

Such rapid time-scales of divergence in the absence of any external input suggest that noise in the NMC-model does not average out. Instead, activity is inherently probabilistic, with a high internally generated variability. Throughout the remainder of this study, we will continue to quantify internally generated variability by the divergence of activity from identical initial conditions.

### Variability is robust across dynamical states

In addition to the *microscopic* divergence of individual somatic membrane voltages, *macroscopic* fluctuations in population spiking activity (**Fig. 2a1**) and population firing rate (**Supplementary Fig. 2**) also diverged rapidly for *t* > *t*_0_. These global fluctuations indicate substantial shared variability between individual neurons. However, the nature of these global fluctuations depends on the balance between excitatory and inhibitory currents (EI-balance) in^30^ the network.

**Figure 2.**
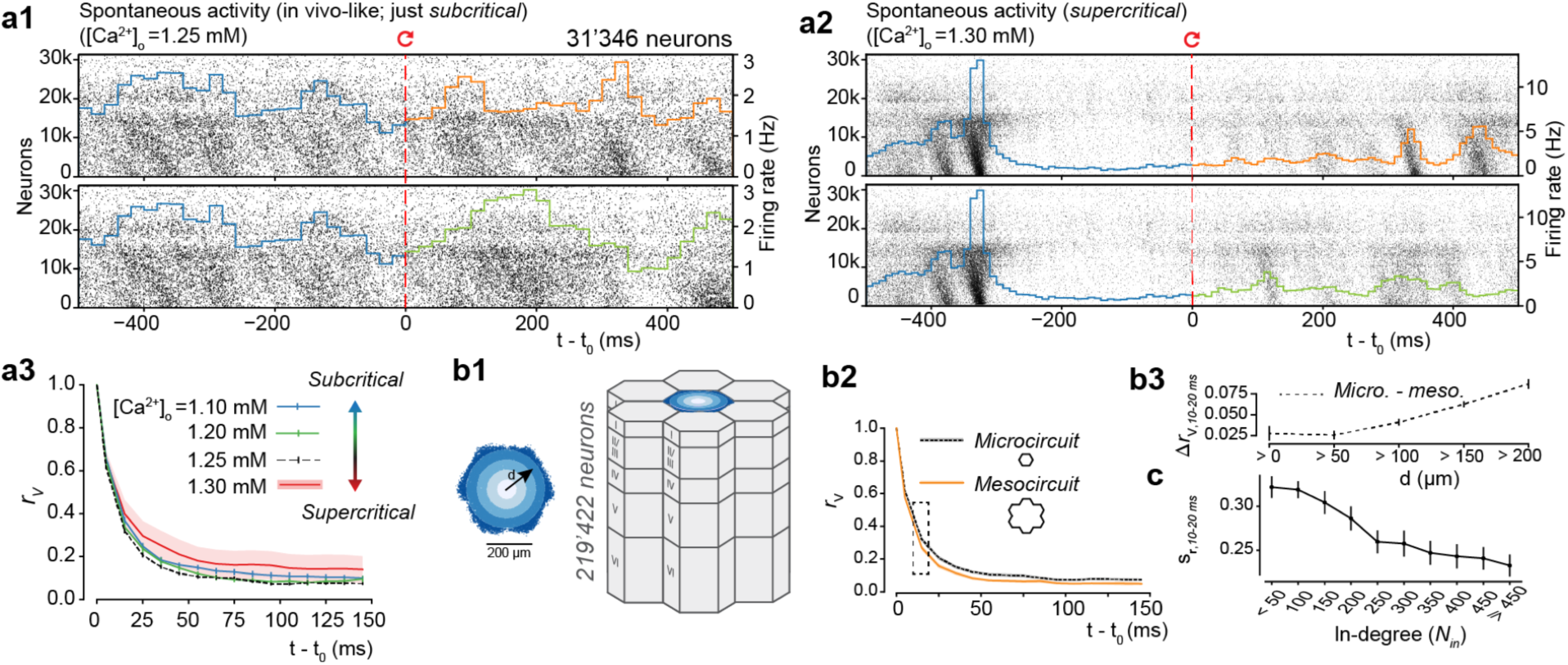
Robust rapid divergence across dynamical states and microcircuit scale. Population raster plot and population peristimulus time histogram (PSTH) for all 31’346 neurons in the microcircuit, during in vivo-like spontaneous activity. Neurons are ordered according to cortical depth, with deep layers at the bottom and upper layers at the top. Each row represents the spikes of one neuron. For visibility, raster lines extend over dozens of rows for each neuron. For *t* < *t*_0_, the top and bottom raster plots show the same simulation, whereas for *t* > *t*_0_, the raster plots depict two simulations resuming from identical initial conditions at *t*_0_, but using different random number seeds. (**a2**) Same as **a**, but for supercritical activity. (**a3**) *RMSD_V_* and *r_v_* across dynamic regimes (20 saved base states, mean ± 95% confidence interval; same as **Figure 1d** for in vivo-like regime ([*ca*^2+^]_*0*_ = 1.25 *mM*).(**b1**) The microcircuit (center, blue), surrounded by 6 other microcircuits (grey), forming a continuous mesocircuit of ∼220’000 neurons, with no boundary effects between the circuits. (**b2**) *r_v_* for the center microcircuit when simulated without surrounding circuits (black), and of the center microcircuit when simulated as a mesocircuit (orange) (microcircuit: 40 saved base states; mesocircuit: 20 saved base states; mean ± 95% confidence interval). (**b3**) Quantifying edge effects. Difference of *r_v_* between the same neurons in the microcircuit and the mesocircuit at 10-20 ms, plotted according to distance from horizontal center (mean ± 95% confidence interval). (**c**) Similarity *s_r_* for subsets of neurons grouped by in-degree (bin size: 50; mean ± 95% confidence interval).

In the NMC-model, the EI-balance is modulated through the effects of extracellular calcium concentration [*ca*^2+^]_*0*_ on vesicle release probabilities. As the dependence on [*ca*^2+^]_*0*_ is stronger for excitatory than for inhibitory synapses, increases in the concentration of [*ca*^2+^]_*0*_ lead to stronger relative excitation and a sharp transition from asynchronous states (*subcritical*) to more correlated activity^17^, which is regenerative and synchronous (*supercritical*; **Fig. 2a2**).

In the in vivo-like state analyzed here [*ca*^2+^]_*0*_ = 1.25 *mM*), the microcircuit is in a just subcritical^31^ state of asynchronous spontaneous activity, where it reproduces several findings from in vivo experiments^17^. While this asynchronous state might be important for efficient coding^32,33^, the exact E-balance in vivo is difficult to determine, and is likely to reconfigure dynamically as a function of the state of arousal and attentiveness of the animal.^34^ We therefore investigated the relationship between the time course of divergence and different dynamic regimes. We observed that the rapid divergence of electrical activity was approximately conserved across these different dynamic states (**Fig. 2a3**). While steady-state electrical activity was slightly more de-correlated in the in vivo-like state, the time course of divergence was remarkably similar. We also found that the synchronous state still displayed high shared variability, with unpredictable timing of population bursts (**Fig. 2a2**, *t* > *t*_0_). In our model, therefore, intrinsic variability, as quantified by the time course of divergence, is conserved across a spectrum of dynamic states and does not depend on the exact EI-balance.

### Variability is nearly saturated at the scale of the microcircuit

It is possible that the amount of internally generated variability depends not just on the dynamic state of the model circuit but also on its size. We have previously shown that in models of the size used in the simulations just described, dynamic states stabilize ^17^. At this size, dendritic trees and thus the afferent connections of neurons in the horizontal center of the microcircuit are fully located within the microcircuit. However, a large fraction of their recurrent connections with neurons in the surrounding tissue are with neurons at the periphery of the microcircuit. Since these were not included in the simulations, large portions of synaptic input to peripheral neurons were missing. To quantify the effect of this additional input on variability in the microcircuit, we surrounded the original microcircuit with six additional microcircuits, simulating a much larger *mesocircuit*, which provided missing synaptic input to the neurons at the periphery of the microcircuit (**Fig. 2b1**, blue and grey). Connectivity in this mesocircuit was homogeneous, both within and between the individual microcircuits.

When we compared the divergence of membrane potentials between micro - and mesocircuit simulations, we found that membrane potentials diverged slightly faster in the mesocircuit, although the time courses of divergence followed similar trends (**Fig. 2b2**). The mean difference in *r_v_*(*t*) was always below 0.06, and the steady state difference below 0.03. We next focused on the difference at 10-20 ms, which we found to be a good predictor of the relative order of differences at any time. We found that *s_r_*_10–20 *ms*_ was directly related to distance from the horizontal center, with the largest differences in neurons at the periphery of the microcircuit (**Fig. 2b3**). At the periphery, the increase in variability between meso- and microcircuit simulations was above 0.08, decreasing toward the center and converging just below 0.03 for neurons within 100 *µm* of the center. This suggests that direct additional synaptic input onto a neuron increases variability, but that this additional synaptic input has a weak effect on indirectly connected neurons whose inputs are already saturated. Thus, at the scale of the microcircuit, the amount of internally generated variability is nearly saturated, while variability for neurons at the periphery is underestimated.

### Highly connected neurons diverge faster

To directly quantify the dependence of the time course of divergence on the amount of the synaptic input, we examined the relationship between the similarity *s_r_*(*t*) of a given neuron and the number of connections it receives from within the microcircuit (*in-degree*). Once more, we found that the time course of divergence was faster, the more synaptic inputs a neuron received, as summarized by *s_r_*(*t*) at 10-20 ms (**Fig. 2c**). Thus, it appears that neurons which are more strongly coupled to the local population^35^ are also more likely to diverge quickly. Repetition of the analysis using *RMSD_v_*(*t*) instead of *r_v_*(*t*) gave qualitatively similar results (data not shown). We note that *RMSD_v_*(*t*) and *r_v_*(*t*) are generally highly correlated (**Supplementary Fig. 3a**, *abcd*). In what follows, we hence present the divergence in terms of *r_v_*(*t*), except when there is a qualitative difference.

### Noise amplified by chaos determines internally generated variability

Thus far, we have demonstrated a high level of variability which is robust across dynamical states and nearly saturated at the scale of the microcircuit. We have also shown that divergence is faster for neurons that are more tightly coupled to the local population (**Fig. 2c**). This suggests that the variability of individual neuron activity is driven by the variability of local population activity, or that additional synaptic input simply adds more synaptic noise, or that the noise is determined by some combination of the two effects. In other words, while cellular noise is the only original source of variability in the NMC-model, the question remains to what degree this noise is amplified by recurrent network connectivity.

To address this question, and more generally, to study the interaction of noise sources and recurrent network dynamics, we performed two complementary sets of simulation experiments. In the first set, we sought insights into the role of network dynamics without noise sources, probing the sensitivity of a completely deterministic version of the model to a weak, momentary perturbation. In the second, we studied the opposite case of variability due to stochastic noise sources without amplification by the network.

To implement the first set of simulations, we disabled stochasticity of cellular noise sources, including synaptic transmission, by using a fixed sequence of random numbers, which made the random outcome deterministic (or alternatively by completely replacing the stochastic model with a deterministic one, see below). This enabled us to observe amplification of perturbations through the network without the effect of continuously varying cellular noise sources. As the sole source of perturbation, we injected a single extra spike into one of the neurons in the microcircuit (see Methods). We observed that the network diverged rapidly (**Fig. 3a1**, dashed line), though more slowly than with noise sources enabled (**Fig. 3a1**, solid line). In fact, even a miniscule current injection, which shifted the majority of spike times by less than 0.05 ms (see Methods), eventually led to a divergence of membrane potentials similar to the divergence observed in the full model with noise sources (**Fig. 3a1**, dotted line). The slightly higher steady-state correlation *r*_∞_ in the deterministic simulation was due to identical spontaneous release of neurotransmitter, identical ion-channel opening probabilities, and the small, but identical, noisy component of the depolarizing current injection. However, the relative difference in *RSMD*_∞_ was much smaller than the difference between the deterministic and the stochastic simulations (**Fig. 3a2**, top vs. bottom). That is, any perturbation to the system eventually led to a similarly large steady-state divergence. We conclude that the underlying dynamics of the circuit are chaotic, in the sense that small perturbations, such as one injected spike, lead to completely different, unpredictable activity.

**Figure 3.**
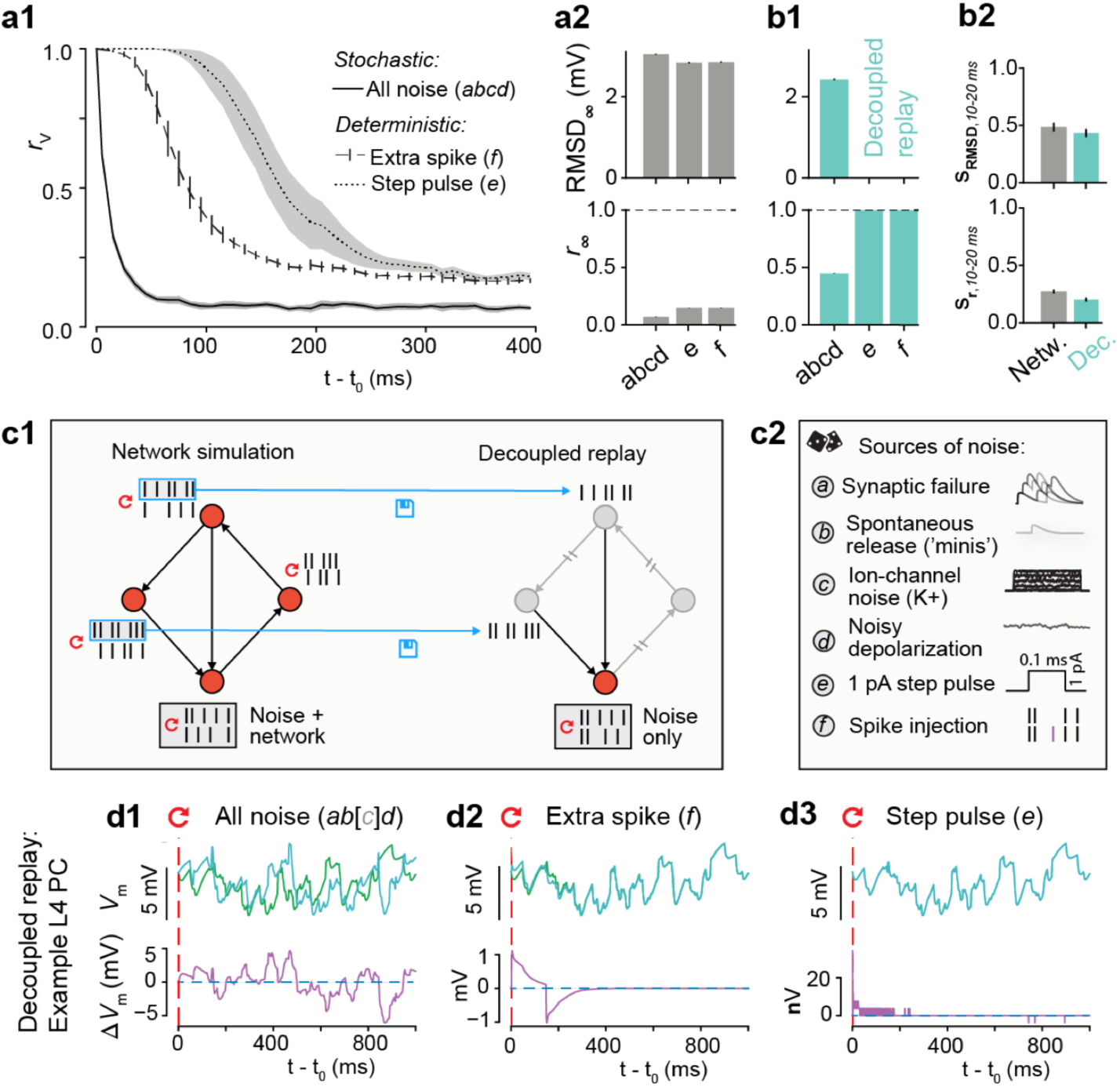
Noise amplified by chaos determines internally generated variability. (**a1**)Time course of correlation *r_v_* after resuming at t_0_ from identical conditions with different forms of perturbation. Full cellular noise as before, solid line (*abcd*); no cellular noise, but perturbing with a single extra spike in one neuron, dashed line (*f*); a miniscule step pulse perturbation in all neurons, dotted line (*e*). (*abcd*: 40 saved base states; *e, f*: 20 saved base states; mean ± 95% confidence interval) (**a2**) Steady-state root-mean square deviation *RMSD*_∞_ and correlation r_∞_ for stochastic (*abcd*) and deterministic simulations (*e, f*) as defined in a1 (mean ± 95% confidence interval). (**b1**) As in **a2**, but for decoupled, replayed simulations. (*b2*) Similarity *s_RMSD_* and *s_r_* at 10-20 *ms* with all noise sources enabled, for network and decoupled simulations (mean ± 95% confidence interval). (**c1**) Decoupled replay paradigm. Presynaptic spike trains from a network simulation are saved and then replayed to the synapses of each neuron in a decoupled simulation, thereby removing variability due to feedback network dynamics. (**c2**) Overview of sources of noise and perturbations. (**d**) Decoupled replay simulations (see **c1**) for a representative L4 PC neuron, with somatic membrane potential differences between the two trials only due to cellular noise sources (*ab[c]d*), a single extra presynaptic spike (*f*) or a miniscule step-pulse perturbation (*e*). [*c*] indicates that for some neuron types in the NMC-model, such as L4 PCs, no stochastic ionchannels are present.

It is important to note that when using a fixed random seed to make the stochastic version of the Tsodyks-Markram synapse model deterministic^17,36^, any extra or missing presynaptic spike can change the outcome for the next spike by advancing the sequence of random numbers. To avoid this difficulty, we ran equivalent simulations using the deterministic version of the Tsodyks-Markram synapse model (see Methods). In these simulations, extra spikes and small perturbations produced qualitatively similar divergence time courses (**Supplementary Fig. 4a** vs. **4b**, dark green and pink lines).

We had shown that the network amplifies extra spikes or even small perturbations of membrane potentials. This leads to chaotic divergence of activity with similar steady-state variability, but different time courses. It remained to be seen whether this high level of variability requires network amplification or whether it could be generated by the noise sources alone.

To address this question, we implemented a second set of simulations to study the case of ongoing noise sources without network propagation. In these *decoupled replay* simulations, in contrast to regular *network simulations*, synaptic mechanisms were activated by spikes at fixed times, recorded in an earlier simulation experiment (**Fig. 3c1**). In this way, the network was no longer able to amplify neuronal variability and neuronal variability was entirely due either to cellular noise sources or perturbations (**Fig. 3d**). We found with all noise sources turned on, somatic membrane potentials still diverged rapidly, as quantified by *s_r_*,_10-20 ms_ (**Fig. 3b2**) (as mentioned above, we found *s_r_* at 10-20 ms to be a good predictor of the relative order of *s_r_* at any time). However, steady-state *r*_∞_ was higher and *RMSD*_∞_ was lower than in the network simulations (**Fig. 3b1** vs **Fig. 3a2**). When the decoupled replay paradigm was used with the deterministic version of the model, single extra spikes and brief current injections only evoked small, transient perturbations (**Fig. 3d2**,3). It follows that the high level of variability observed in network simulations was due to chaotic network dynamics which amplified rapid perturbations of activity from cellular noise sources.

### Synaptic noise dominates variability

To understand the contribution of individual noise sources in this interplay of noise and recurrent network dynamics, we designed a series of simulation experiments where we selectively disabled specific subsets of noise sources, instead of all of them as in the deterministic version above. We observed that disabling all noise sources except synaptic failure produced a time course for *r_v_*(*t*) and steady-state divergence *r*_∞_ which was very similar to observations with all noise sources combined (**Fig. 4a1**, black and green lines). On the other hand, disabling all but ion channel noise or all but the noisy current injection led to much slower divergence (**Fig. 4a1**, orange and purple lines). As before, we quantified the speed of divergence by the similarity *s*_*r at 10-20 ms*_ after *t_0_* (*s_r_*,10-20*ms* (**Fig. 4a3**, cyan). Our results suggest that simulations with synaptic failure give rise to rapid divergence, whereas steady-state *r*_∞_ and *RMSD*_∞_ depend on noise sources only weakly (**Fig. 4a2**). We conclude that in the NMC-model, the time course of divergence depends on synaptic noise, a combination of synaptic failure and spontaneous release, and that other noise sources add little to no additional variability.

**Figure 4.**
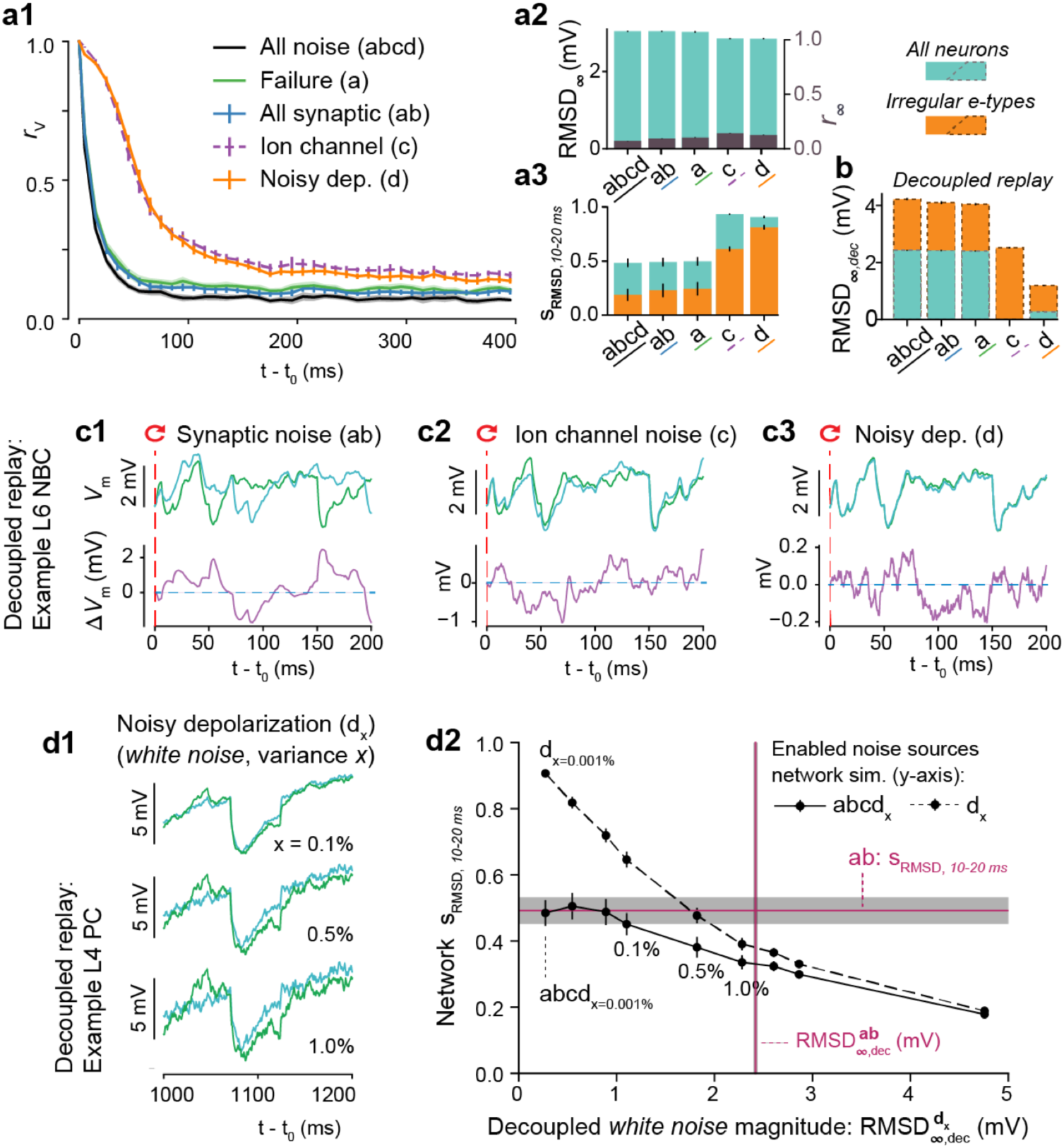
Synaptic noise dominates variability. (**a1**)Time course of correlation *r_v_* after resuming at t_0_ from identical conditions with different noise sources enabled (*abcd*: 40 bases states; a, ab, c, d: 20 base states; mean ± 95% confidence interval). (**a2**) Steady-state root-mean square deviation *RMSD*_∞_ (cyan) and correlation *r*_∞_ (purple) with different noise sources enabled. (**a3**) Similarity *s_RMSD_* at 10-20 ms with different noise sources enabled, for all neurons (cyan) and irregular e-types (orange). (**b**) Steady-state root-mean square deviation for decoupled simulations, *RMSD*_∞,*dec*_, for all neurons (cyan) and irregular e-types (orange). Only irregular e-types in (c), 1,137 out of 31,346 neurons. (**c**) Decoupled replay simulations for a representative L6 NBC neuron, with somatic membrane potential differences between the two trials only due to synaptic noise (*ab*), ion-channel noise (*c*) or a noisy current injection (*d*). (*d1*) The effect of changing random seeds for the noisy depolarization only, for different noise strengths in a decoupled simulation. x: white noise variance as percentage of mean injected current (**d2**) The decoupled steady-state membrane potential fluctuations 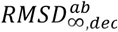, evoked by different magnitudes of white noise without network dynamics, versus the similarity *s_RMSD_* at 10-20 ms during network simulations when either turning on only the white noise depolarization (*d*) or all noise sources (*abcd*). Similarly, in purple, 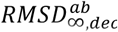 for synaptic noise versus the similarity at 10-20 *ms* when only turning on synaptic noise (*ab*). All error bars and shaded areas indicate 95% confidence intervals. Means for **d2** are based on ten base states.

### Ion-channel noise in irregular firing neurons is overshadowed by synaptic noise

Synaptic noise in the NMC-model is modeled at every single synapse, while ion-channel noise is limited to irregular firing e-types ^17,19^. Irregular e-types are defined by high intrinsic spike-time variability in response to constant current injections in vitro, even in the absence of synaptic noise. In the NMC-model, irregular spiking is modeled with a subset of stochastic ion-channels, in accordance with in vitro findings on the source of the irregular spiking patterns observed in cortical interneurons^20^. In contrast, regular firing e-types do not require noisy ion-channels to replicate in vitro spiking behavior. To better understand the interplay of ion-channel noise and synaptic noise, we focused our next analysis solely on irregular firing e-types. We observed that irregular firing e-types diverged significantly faster than the whole population (**Fig. 4a3**, orange vs. cyan). However, synaptic noise still dominated over ion-channel noise. Enabling ion-channel noise in addition to synaptic noise led to only marginal gains in divergence rate; when ion-channel noise was enabled on its own, divergence was significantly slower (**Fig. 4a3**, orange, *ab* vs. *abcd* and *c*). This suggests that in in vivo conditions, noise from stochastic ion-channels is probably overshadowed by synaptic noise. This contrasts with in vitro conditions, where channel noise is the only major noise source.

### Synaptic noise acts as threshold for other noise sources

There are in reality many smaller noise sources that are not included in our model (see Discussion). To understand how additional noise sources of various magnitudes could influence divergence, we analyzed the magnitude of the previously analyzed cellular noise combinations in a decoupled replay, with network propagation removed (*RMSD*_∞,*dec*_) (**Fig. 4b**; see Figure 4c1-3 for representative examples). We found that this magnitude inversely relates to the rate of divergence, *s*_*RMSD*,10-20 *ms*_ (**Fig. 4a3**). That is, a larger *RMSD*_∞,*dec*_ leads to a faster divergence (as measured by a smaller *s_RMSD_,*_10-20 *ms*_) (see also **Supplementary Fig. 5** for an extensive comparison of noise sources across simulation paradigms). In the NMC-model, synaptic noise has the largest *RMSD*_∞,*dec*_ and determines the rate of divergence. But how strong would any other noise source have to be to generate network variability that is detectable beyond synaptic noise? To answer this question, we studied how the magnitude of an unknown noise source affects the time course of divergence. As a proxy for unknown noise sources, we increased the variance 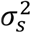 of the injected white noise depolarizing current. Previously, the variance had been set to 0.001% of the firing threshold for each neuron—a level far lower than other sources of noise. When we increased the variance to values from 0.01% up to 10% and disabled all other noise sources, we observed that increasing 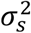 led to more rapidly diverging network dynamics (**Supplementary Fig. 6a**). However, when other noise sources were also enabled, the noisy current injection only affected network dynamics beyond a certain threshold (**Supplementary Fig. 6b**).

To characterize this threshold, we first measured 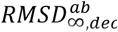, that is, the steady-state divergence of membrane potential fluctuations evoked by noisy current injection alone in a decoupled replay, for various levels of 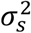 (**Fig. 4d1**). We then compared 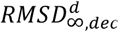 to the time course of divergence in the corresponding network simulations with the same noise conditions (i.e. only noisy depolarization (d); **Fig. 4d2**, dashed line). We found that the rate of divergence as measured by *s_r_*,_10-20 *ms*_ was strongly dependent on 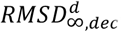, with larger values leading to faster divergence. In contrast, when we repeated the analysis with *all* noise sources enabled (**Fig. 4d2**, solid line), the dependence on 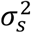 was weaker, indicating a smaller impact of 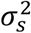 on *s_r_*,_10-20 *ms*_. Indeed, 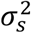 only had a meaningful influence when it was beyond a threshold in the range 0.1% −0.5%. At this threshold, the steady-state divergence in decoupled replays 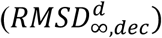 evoked by the noisy current alone was just above 1 mV, approximately half of the value for synaptic noise sources 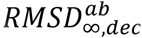, **Fig. 4d2**, vertical purple line ^“^*ab*^”^). When 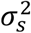 is increased even more, the curves for *s_r_* _10-20 *ms*_ with noisy current alone and with all noise sources eventually began to converge. Thus, when 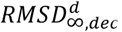 was larger than 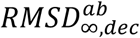 the noisy current injection dominated other noise sources. This suggests that the strongest source of cellular noise dominates over other sources, unless they are of a comparable magnitude. Under biological conditions, we predict that synaptic noise dominates. This prediction matches previous findings that cortical neurons respond very reliably to current injections in vitro (no synaptic noise)^12^. However, it also suggests an entirely different picture of the reliability of neuronal responses to presynaptic inputs in vivo, with synaptic noise contributing to variability.

### Low trial-by-trial spike-timing reliability during spontaneous activity

Neurons transmit signals to postsynaptic partners only in the form of spikes. Their timing is determined by a non-linear transformation of the somatic membrane potential (*V_m_*) we analyzed so far. We therefore next characterized the role of the spike generation mechanism in influencing the reliability of neural responses, and compared both the variability of membrane potentials and spike times between independent network simulations of spontaneous activity over 30 independent trials with different initial conditions. We found that membrane potentials (**Fig. 5a**, top) and the corresponding spike trains (**Fig. 5b**, top) were both highly variable. We then used the spike times recorded from each of these network trials in five decoupled replay simulations per trial. As before, we observed that membrane potentials were less variable and more correlated in decoupled simulations (**Fig. 5a**, bottom). Indeed, the distributions of *r_v_* in decoupled and network simulations were almost completely disjoint (**Fig. 5c1**), with decoupled replay simulations exhibiting much more correlated membrane potentials overall. However, considering just the spike times, we found no clear difference in variability between the network (**Fig. 5b**, top) and the decoupled replay simulations (**Fig. 5b**, bottom). In stark contrast to the results for membrane potentials, quantification of spike time variability using a correlation-based measure, *r_spike_*^37^, showed a large overlap in the distributions for decoupled and network simulations (**Fig. 5c2**, red area vs. solid black line; *σ_spike_* = 5 ms), in stark contrast to the case for membrane potentials. This suggests that the spike initiation mechanisms cannot transform the increased reliability of *V_m_* into reliable spike trains during spontaneous activity. Indeed, we found that an increase in the magnitude of *r_v_* did not predict a corresponding increase in *r_spike_* (**Fig. 5c3**). On the contrary, the two measures displayed a weak inverse correlation. We note that we found no decrease in variability when other sources of noise besides synaptic noise were disabled (**Fig. 5c2**, dashed brown line), as expected in light of our previous result that synaptic noise accounts for a large proportion of variability.

**Figure 5.**
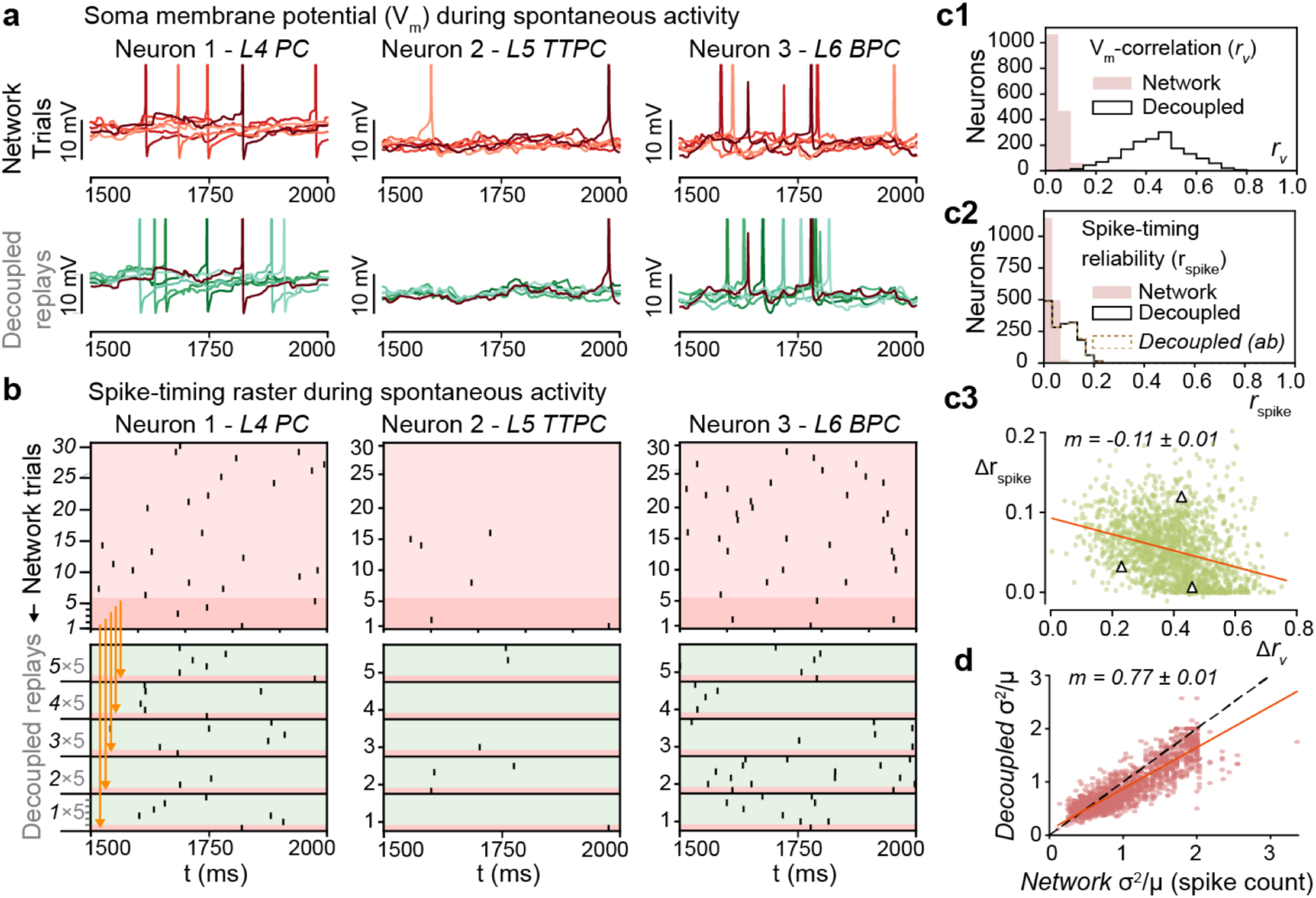
Low trial-by-trial spike-timing reliability during spontaneous activity. (**a**)Somatic membrane potentials (*V_m_*) of three representative neurons. Top: during six independent trials of spontaneous activity. Bottom: five decoupled replay trials (green) with the same presynaptic input as during the original network simulation trial (red), but with different random seeds. (**b**) Top: Raster plot of spike times for the same example neurons as in A, during 30 independent trials of spontaneous activity. Bottom: 5 decoupled replay trials (green) of the same input received during 5 of the 30 original trials (dark red). (**c1**) Mean somatic membrane potential correlation *r_v_* of the 1666 (*ab*: 1670) most central (and spiking) pyramidal neurons from layers 4, 5, and 6 between independent network simulations, and between decoupled replay simulations with identical presynaptic inputs. (**c2**) Mean rspike-timing reliability *r_spike_* of the same neurons. Decoupled and decoupled (ab) are overlapping. (**c3**) Change in correlation, ∆*r_v_*, versus change in spike-timing reliability, ∆*r_spike_*, for each neuron for decoupled replay simulations relative to network simulations (linear fit with 68% confidence interval on slope m, red line). Triangles indicate values of representative neurons in panel B. (**d**) Comparison of variance of spike count between network and decoupled replay simulations (same neurons as in **c-e**; linear fit as in **c3**, red line; identity line, black dashed line).

It is possible of course that spike time reliability within tens of milliseconds could be too restrictive a measure of spiking reliability. Therefore, we also compared the variability of spike counts across the entire 0.5 s window analyzed. Use of identical presynaptic inputs produced only a marginal reduction in the variance of spike counts (**Fig. 5d**). In brief, the reliability of spike generation across time-scales is directly, and severely constrained by synaptic noise, even without amplification through network dynamics.

### Rapid divergence of evoked, reliable activity

In the NMC-model, thalamic inputs can evoke responses with varying degrees of reliability among trials^17,38^. What then are the roles of synaptic noise and chaotic network dynamics during these evoked responses? To answer this question, we simulated electrical activity in response to a naturalistic thalamocortical stimulus (**Fig. 6a1**), consisting of spike trains recorded in the ventral posteromedial nucleus (VPM) during sandpaper-induced whisker deflection in vivo^39^. These spike trains were then applied to different feed-forward VPM fibers in the model to achieve a biologically-inspired, time-varying synchronicity among inputs (**Fig. 6a3**; see Methods; see Reimann *et al*.^38^). To avoid introducing external variability on top of the intrinsically generated microcircuit variability, presynaptic inputs were kept identical across trials, but with thalamocortical synapses subject to the same synaptic noise as cortical synapses. The thalamocortical presynaptic inputs were not subject to recurrent network dynamics. Since this condition excludes variability in the system up to and including the thalamus, it can be considered an intermediate stage between the decoupled replay and regular network simulations. The simulations allowed us to identify an upper bound on the reliability of thalamocortical responses. Mean *r_v_*(*t*) during evoked activity was stronger than during spontaneous activity, moving between ∼0.1 and ∼0.4 (**Fig. 6a2**), confirming that that the responses of neuron membrane potentials to the stimulus were relatively more reliable across trials.

**Figure 6.**
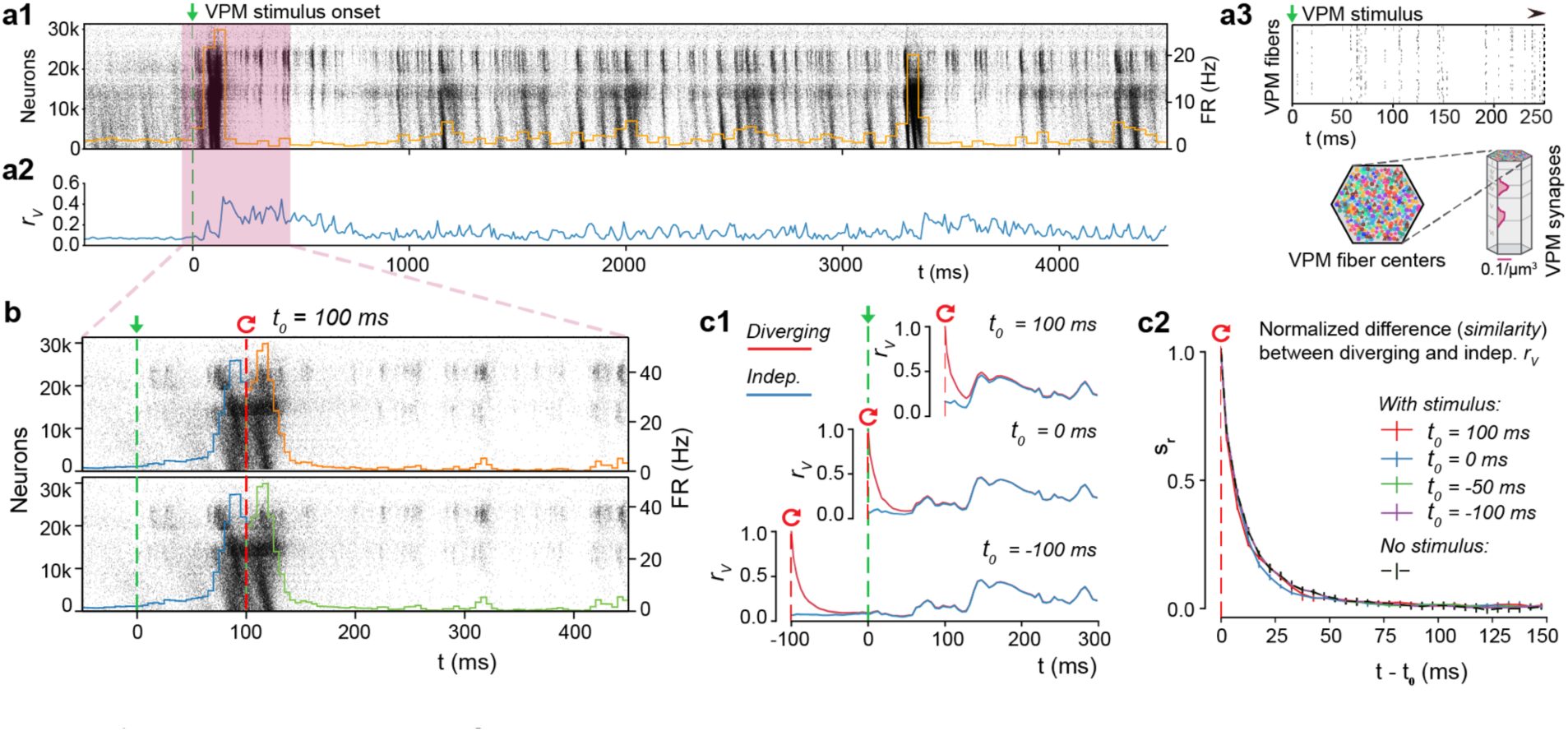
Rapid divergence of evoked, reliable activity. (**a1**)Population raster plot and population peristimulus time histogram (PSTH) for all 31’346 neurons in the microcircuit, during evoked activity with a thalamic (VPM) stimulus. Neurons are ordered according to cortical depth, with deep layers at the bottom and upper layers at the top, and each row representing the spikes of one neuron. For visibility, raster lines extend over dozens of rows for each neuron. (**a2**) Mean somatic membrane potential correlation *r_v_* between independent simulations of the same VPM stimulus (mean ± 95% confidence interval). (**a3**) Schematic of the VPM stimulus. Top: Raster plot spike times for the first 250 ms of the thalamic stimulus. Bottom: 310 VPM fiber centers are assigned 30 colors, and those with identical colors are provided with duplicate spike trains. The synapse density profile across layers for each fiber is shown to the right. (**b**) For *t* < 100, the top and bottom raster plots show the same simulation, whereas for *t* > 100, the raster plots depict two resumed simulations starting from the same saved state at *t*_0_ = 100, using different random number seeds. (**c1**) Resuming from identical initial conditions at different times: during (top), at onset (middle), or before the stimulus (bottom). Mean *r_v_* between independent simulations (blue, as in **a2**), and mean *r_v_* between simulations starting from the same base state (red; mean ± 95% confidence interval). (**c2**) The similarity, *s_r_*, defined as the difference between the *r_v_* of diverging and independent trials, normalized to lie between 1 (identical) and 0 (fully diverged) (mean ± 95% confidence interval). Means are based on 20 base states, *no stimulus* (spontaneous activity) on 40 as before.

To characterize the nature of chaotic network dynamics during this evoked, reliable activity, we again resumed from identical initial conditions, with *t*_0_ at various times relative to the stimulus onset at t = 0 ms (**Fig. 6b**, for *t*_0_ = 100 ms). The population spiking activity across pairs of trials after resuming appeared almost identical, even for time intervals much larger than the divergence time characterized above (**Fig. 6b**). At first glance, it would appear that the input had fully overcome the chaotic divergence. However, quantification of variability by time course of divergence of membrane potentials, *r_v_*(*t*), showed that it dropped rapidly towards the independent trial average (**Fig. 6c1**, top). When we resumed from identical initial conditions at different times, for example at the onset of evoked activity (**Fig. 6c1**, middle) or before onset (**Fig. 6c1**, bottom), *r_v_*(*t*) dropped in the same way, subsequently converging to the average for independent trials. Indeed, *s_r_*(*t*), the normalized difference between the resumed and independent *r_v_*(*t*) showed a pattern of divergence remarkably similar to the divergence observed in simulations of spontaneous activity (**Fig. 6c2**). Resuming from a base state at the peak of evoked activity, *s_RMSD_*(*t*) drops even faster (**Supplementary Fig. 7a**). A simpler stimulus, designed to imitate a whisker flick-type experiment ^17^, yielded comparable results (**Supplementary Fig. 7b,c**). Hence, any neuronal activity, whether spontaneous and unpredictable, or evoked and reliable, is ultimately constrained by similar chaotic network dynamics.

### Spike-timing reliability amid noise and chaos

At first glance, our observations of reliable population spike responses and chaotic divergence of membrane potentials seem to be mutually exclusive. Could it be that membrane potential reliability is simply not correlated with spike-timing reliability, as we observed for the case of spontaneous activity? To answer this question, we again compared network simulations with decoupled replay simulations, which have no network propagation of activity (**Fig. 7a3**). As before, *r_v_*(*t*) was much larger in the decoupled simulations (**Fig. 7a1**, black) than in the network simulations (**Fig. 7a1**, red; same as **Figure 6a2**). However, the difference between the two was always smaller during evoked activity (**Fig. 7a2**, after 0 ms) than during spontaneous activity (**Fig. 7a2**, before 0 ms). This suggests that network dynamics play a reduced role in generating variability during evoked activity. When we focus on individual neurons (**Fig. 7b**), we can see that that the difference between network and decoupled *r_v_*(*t*) at times collapses to zero (**Fig. 7c**). In other words, variability due to network dynamics can intermittently be completely overcome for a sub-population of neurons in the network. Looking at the corresponding membrane potential traces, we observe that these moments occur during periods of reliable spiking (**Fig. 7b**). During evoked activity, in contrast to spontaneous activity, moments of reliable membrane potentials can translate into reliable spiking, at least for some neurons.

**Figure 7.**
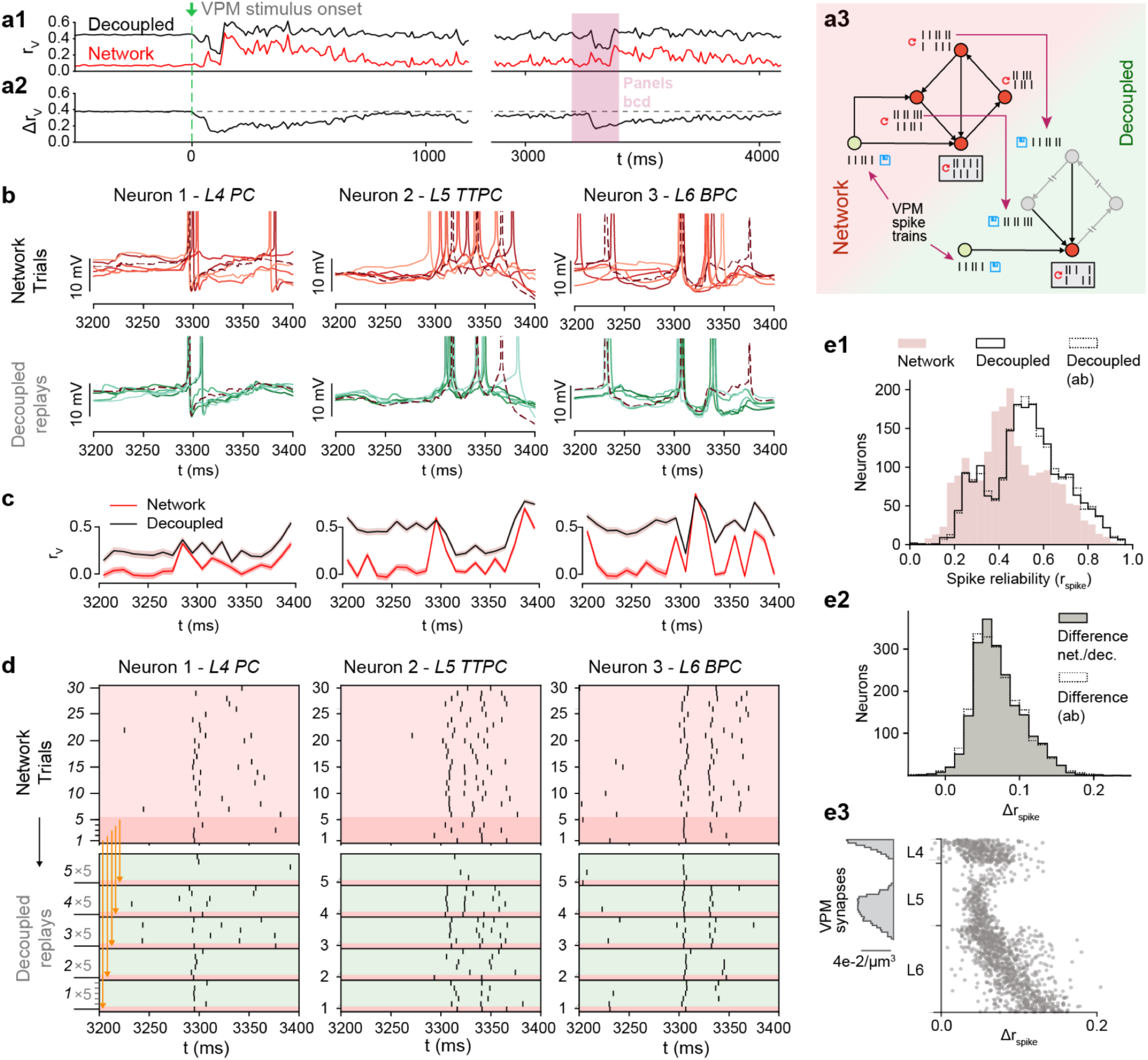
Spike-timing reliability amid noise and chaos. (**a1**) Mean somatic membrane potential correlation, *r_v_*, between independent simulations, and between decoupled replays of those simulations (network simulation identical to **Figure 7A2**). **a2**) Difference in *r_v_* for decoupled and network simulations. **a3**) Schematic of network and decoupled replay simulation paradigms, including thalamic input. **b**) Somatic membrane potentials (*V_m_*) of three representative neurons for the time interval highlighted by the red box in **a**. Top: during six independent trials. Bottom: five decoupled replay trials (green) with the same presynaptic input as during the original network simulation trial (red), but with different random seeds. **c**) Network and decoupled *r_v_* as in a, but only for the three sample neurons in **b**. (**d**) Top: Raster plot of spike times of the same three example neurons as in b, during 30 independent trials of evoked activity. Bottom: Decoupled replay trials (green) of the same input received during 5 of the 30 original trials (dark red). **e1**) Mean spike-timing reliability *r_spike_* of 2024 pyramidal neurons from layers 4, 5, and 6 between independent network simulations, and between decoupled replay simulations with identical presynaptic inputs. **e2**) Difference between *r_Spike_* of decoupled and replayed simulations. **e3**) Difference between *r_spike_* of decoupled and replayed simulations versus position of somata across layers 4,5 and 6 of microcircuit (1675 neurons).

To get an idea of this effect at the population level, we compared spike time reliability *r_spike_* with and without network dynamics (**Fig. 7d**). We observed that removing network dynamics only moderately increased spike-timing reliability (**Fig. 7e1**, red vs solid black line). In fact, increases in reliability were small for all neurons (**Fig. 7e2**, solid black line). In stark contrast to the spontaneous case, a small population of neurons in the network simulations achieved close to perfect spike reliabilities (**Fig. 7e1**). As expected, most of the noise effects could be explained by synaptic noise alone (**Fig. 7e1**,2, dotted black line). We conclude that external stimuli can sparsely and transiently overcome chaotic network dynamics for sub-populations of neurons, though with a substantial residual variability caused by synaptic noise (albeit much smaller than during spontaneous activity).

### High reliability requires recurrent cortical connectivity

It is conceivable that the spike-timing reliability we observed could simply be a result of direct and feed-forward input from VPN^40^. Indeed, when we look at changes in reliability without network dynamics, the strongest increase in reliability is in neurons at the bottom of layer six that receive comparatively little direct VPM input (**Fig. 7e3**). On the other hand, the VPM input was weak compared to the recurrent connectivity, making up only 7% of the connections onto neurons in layer 4, 4% for layer 5, and less than 3% for layer 6. To test whether the intermittent suppression of chaotic dynamics is simply an effect of the feed-forward input, we designed a new simulation paradigm similar to our previous decoupled replay, where each neuron received a combination of replayed presynaptic inputs from a simulation of spontaneous activity and from the direct feed-forward VPM input it received in the evoked network simulations (**Fig. 8a1**). That is, each neuron receives input as in a spontaneous activity trial through its recurrent synaptic contacts, and input as in an evoked trial through its feed-forward synaptic contacts.

**Figure 8.**
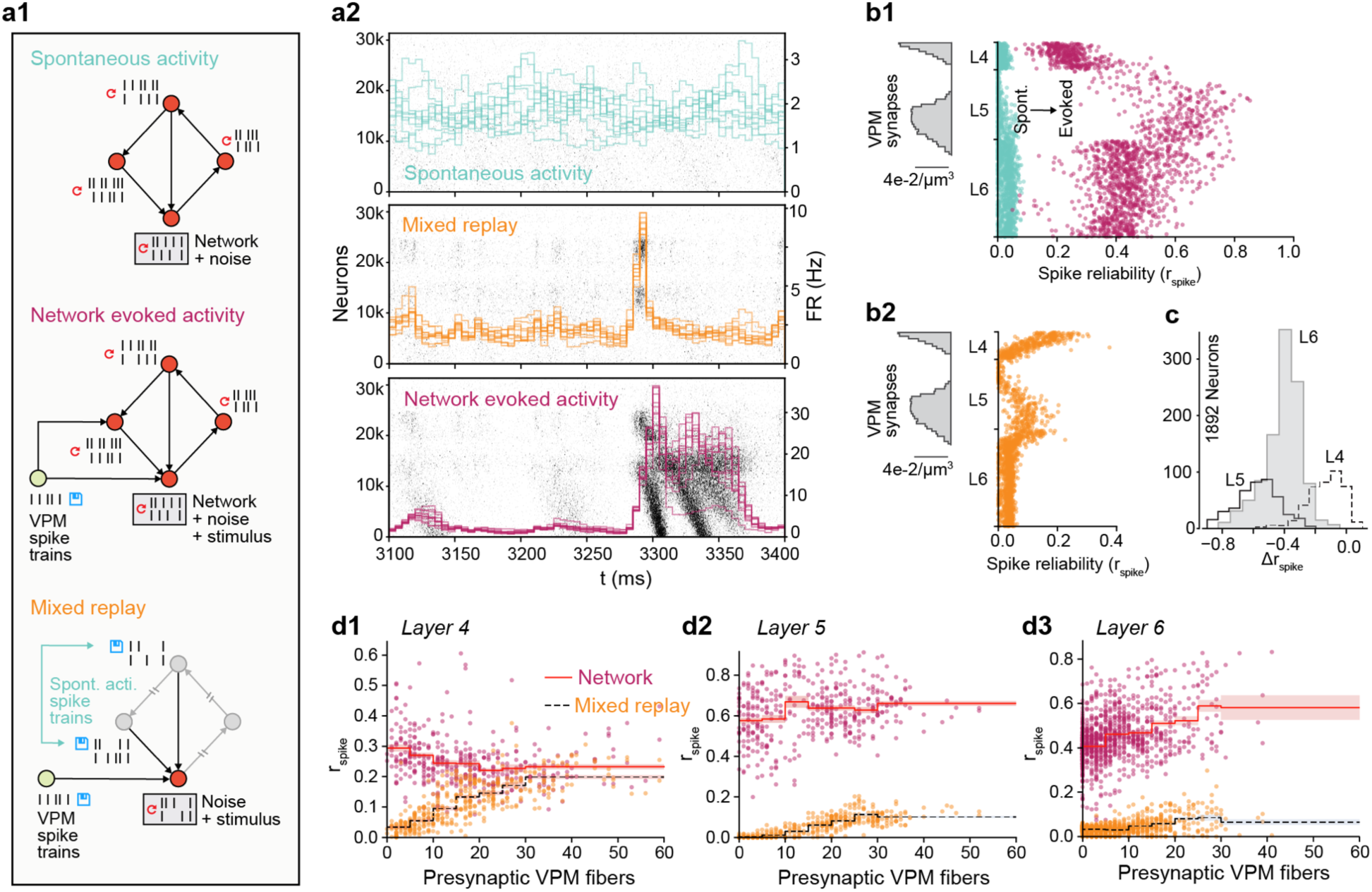
High reliability requires recurrent cortical connectivity. **a1**) Overview of three simulation paradigms: *spontaneous activity, network evoked activity* (with network propagation intact and VPM input), and mixed replay (with network propagation replaced by replays of spontaneous activity spike trains, and VPM input) **a2**) Examples of population spiking activity during the three simulation paradigms. **b1**) Spike-timing reliability, *r_spike_*, during spontaneous (blue) and evoked (purple) activity for 1675 excitatory neurons in the center of layers 4, 5 and 6. **b2**) Spike-timing reliability, *r_spike_*, during a mixed replay with VPM input but with network propagation disabled for the same neurons as in b1. (c) Difference in *rs_Pike_* between evoked activity with and without network propagation for 1892 excitatory neurons in the center of layers 4, 5 and 6 (same for d1-3). (d1) The number of presynaptic VPM fibers from which each neuron receives input versus *r_spike_* in evoked simulations with (network) and without (mixed replay) network propagation.

In this *mixed replay* paradigm, the population response was much weaker (**Fig. 8a2**). While in simulations of evoked activity all neurons showed higher reliability than in simulations of spontaneous activity (**Fig. 8b1**), in the mixed replay, the only cells that showed increased reliability were those close to the VPM synapses (**Fig. 8b2**). Furthermore, the only neurons to display similar reliability, with and without recurrent network propagation, were a small group in layer 4 (**Fig. 8c**). Taken together, these findings suggest that feed-forward VPM input alone is not enough to make the majority of neurons spike reliably.

To test this hypothesis, we compared the reliability between the two simulation paradigms to the number of presynaptic VPM fibers innervating each neuron (Fig. 8d1-3). We can see that neurons in layer 4 that receive little direct VPM input responded more reliably with the network enabled than neurons that receive a lot of VPM input with no network effect (**Fig. 8d1**). Similarly, neurons in layers 5 and 6 were more reliable in mixed replay when they had more presynaptic VPM connections. However, this reliability increases drastically when network dynamics are enabled (**Fig. 8d2**,3). We conclude that the reliable spiking observed in response to VPM inputs is enabled and propagated by recurrent cortical connectivity, and that this is true both for neurons that receive large direct VPM input, and for neurons that receive little or no such input. In brief, in spontaneous activity, recurrent connectivity amplifies variability; in the evoked state, it amplifies reliability.

## Discussion

In the present study, we used a biologically constrained model of a prototypical neocortical microcircuit^17^ to estimate the intrinsic variability of local neocortical activity (Figs. 1-5) and explore the implications for reliable stimulus encoding (Figs. 6-8). We found that cortical circuitry supports millisecond-precision spike-time reliability amid highly variable, chaotic network activity. This resolves a long-standing question: Is the cortex too noisy for the precise timing of a spike to matter^8,12,22,24^? Put simply, if spiking is unreliable, information must be coded by firing rates estimated in populations of neurons^22,24^, whereas if it is reliable, precise spike timing of single neurons could contain significant information^8,12^. Here, we demonstrated cortical circuitry naturally supports both regimes.

This debate has raged on for decades, because the experimental manipulations required to untangle the noise sources in the brain, and evaluate their impact on spike reliability, are impossible to perform in vitro or in vivo. Using the NMC-model, we were able to perform a series of simulation-based manipulations where we systematically added and removed noise sources to quantify their impacts. These manipulations yielded several novel insights.

First, we found that spontaneous activity in cortical circuitry is intrinsically variable, both at the single neuron and population level (Figs. 1,2). While some of the effects of cellular noise sources on variability had been studied in single biophysical Hodgkin-Huxley type neuron models^20,40–42^, this is the first estimate of internally generated variability in an integrated, biologically constrained model of a cortical circuit. Our results confirm previous predictions of simplified network models that showed that biological details such as distance dependent connectivity^43^, feedback inhibition^44^, and differences in synaptic time scales^45^—all intrinsically part of our model—can lead to internally generated variability.

Our second insight was that stochastic synaptic transmission is amplified by chaotic network dynamics to drive a rapid, chaotic divergence of the network, resulting in the above-mentioned variability (Figs. 3,4). Chaotic network dynamics without synaptic noise have been extensively studied^22–24^, and it has been suggested that synaptic noise generates high neural variability in postsynaptic neurons^18,46^. However, this is the first time that the interplay between stochastic synaptic transmission and chaotic network dynamics has been seen and understood.

A third insight was that spike times were unreliable during spontaneous activity (**Fig. 5**), but became reliable during evoked activity (Figs. 6,7). Even comparatively weak thalamocortical input could switch the network to a highly reliable spiking regime. Left alone, the network is in a chaotic regime, but transient inputs can push the network towards a temporally precise regime where millisecond-precision spike-time reliability is possible. This explains how patterns of activity generated by cortical circuitry in response to sensory stimuli can often have millisecond spike-timing precision^26,47^.

The fourth—and perhaps most surprising insight—is the mechanism for this dichotomous behavior. We determined that the recurrent network architecture causes both the amplification of synaptic noise during spontaneous activity, and the quenching of the noise sources in the presence of input. In the former case, the network is driven towards a chaotic, divergent regime, whereas in the latter, a temporally precise regime emerges (**Fig. 8**). The critical role of the recurrent network stands in contrast to previous modelling work which showed that relatively few synchronous thalamic inputs maximize reliability in single neurons in cat visual cortex^40^. However, this study likely overestimated synaptic reliability—synaptic release probabilities are lower in vivo than in vitro, both in general^16^ and in this specific pathway^48^. We conclude that the thalamic input is the trigger, but that the input can only pull the neurons out of chaos with help by the network.

The exact mechanism for this triggering of reliable spiking, and the means by which signals are reliably propagated through the circuitry amid variable activity remains a subject for future investigation. One possible explanation is that certain connectivity motifs could amplify reliability through redundant connectivity. Candidate motifs have already been identified in the NMC-model, such as common neighbor motifs^49^ and high-dimensional cliques that shape spike correlations between neurons^38^. Dendritic nonlinearities, such as N-Methyl-D-aspartate (NMDA)-mediated plateau potentials evoked by clustered synaptic inputs onto the dendritic tree could also play an important role^50,51^.

### Potential effects of missing biological detail

While the NMC-model is one of the most detailed models of neocortical circuitry to date, several biological details are lacking. In terms of noise sources, the most important missing detail is ion-channel noise. Other electrical noise sources such as thermal noise are orders of magnitude smaller^13^.

The partial ion-channel noise in irregular firing neurons in the NMC-model (which is responsible for the irregular initiation of action potentials in vitro^20^) is overshadowed by synaptic noise under in vivo-like conditions (**Fig. 4**). But how would additional ion-channel noise in axons and dendrites of all neurons impact variability? In dendrites, ion-channel noise is thought to evoke little to no variability in isolated back-propagating action potentials^41^. Thus, mean ion-channel models are likely sufficient for accurate action potential initiation.

Action potentials reliably invade axonal arbors of neocortical pyramidal neurons without failures^52^. But as action potentials propagate along axons, their timing becomes increasingly variable. Simulations predict that ion-channel noise affects action potential timing in all axons with a diameter below 0.05 µm, with the standard deviation of action potential variability predicted to increase by 0.6 ms per 2 mm in 0.02 µm diameter axons^21^. In the NMC-model, axons have a mean axonal diameter of around 0.03 µm and are modeled deterministically. Therefore, ion-channel noise in longer axons could increase variability of spike timing by up to several milliseconds.

The missing ion-channel noise might push the circuit towards a more variable state. On the other hand, adding missing detail to the synapse models might increase reliability: The reliability of synaptic transmission increases with the number of readily releasable vesicles^53^. Some studies have found *univesicular* synaptic transmission at cortical synapses^54^, while others have estimated there may be as many as ten releasable vesicles per synapse^55^.

The current version of the NMC-model assumes one readily releasable vesicle per synapse, and thus potentially underestimates synaptic reliability. To estimate the potential impact of *multivesicular* release, we repeated the simulation experiments with an increasing number of readily releasable vesicles (*n_rrp_*) at all synapses (**Supplementary Fig. 8**). As expected, the time course of divergence slowed with increasing *n_rrp_*. Nonetheless, for mean *n_rrp_* values which reproduce cortical PSP variability data (*n_rrp_* = 2 – 3; data not shown), synaptic noise remains the dominant source of noise driving the rapid chaotic divergence. In addition, *n_rrp_* may vary between and across synapse types, but a systematic exploration thereof is beyond the scope of the present study.

There are other intrinsic mechanisms not yet included in the NMC model such as gap junctions, intra-circuit neuromodulation^56^ or active information transfer from glia to neurons^57^,^58^, whose contributions to variability within cortical circuits are as yet poorly understood. However, for these mechanisms to contribute significantly as additional noise sources above and beyond synaptic noise, they would have to cause somatic membrane potential differences on the order of 1 mV (*RMSD*) (**Fig. 4**).

### Concluding remarks

In the intact animal, a neocortical microcircuit is integrated with the rest of the brain and constantly receiving input: around 80% of corticocortical synapses are formed with non-local neurons^17^, which are not yet accounted for in the NMC-model. In the behaving brain, most of this external input to the microcircuit will likely contain signals: for example, visual cortex is strongly modulated by movement-related activity^59^.

This study provides, for the first time, a data-constrained biophysical framework towards theories of cortical coding that can integrate these signals along a spectrum: from population firing rates to reliable individual spike-times. We hypothesize that population firing rates might encode attentional states, general movement-related activity, or other slow variables, whereas patterns of spikes with high temporal precision^26^ might encode more particular information, such as the touch of a whisker or, perhaps, perception of a specific object. The critical role of the recurrent network for the reliable representation of information in these spike patterns further suggests that such patterns might play an important role in computations across the hierarchy of cortical regions^60^. The present study provides a solid foundation for future studies in this direction, and ultimately towards a deeper understanding of cortical information processing.

## Methods

### Simulation

#### Model of neocortical microcircuitry (NMC)

Simulations of electrical activity were performed on a previously published model of a neocortical microcircuit in two-week old rat. Reconstruction and simulation methods are described extensively by Markram *et al*.^17^. In our study, we used a microcircuit consisting of 31,346 biophysical Hodgkin-Huxley NEURON models and around 7.8 million connections forming roughly 36.4 million synapses. Synaptic connectivity between 55 distinct morphological types of neurons (*m-types*) was predicted algorithmically by integrating anatomical data, such as layer-dependent cell type densities, morphologies and bouton densities, to generate a wiring diagram^61^ with highly heterogeneous connectivity^38,62,63^. Consequently, the NMC-model exhibits naturally emerging structural and functional EI-balance^62^, without relying on assumptions about the exact level of coupling between excitatory and inhibitory currents. The densities of ion-channels on morphologically-detailed neuron models were optimized to reproduce the behavior of different electrical neuron types (*e-types*) as recorded in vitro^64^. We also used a larger mesocircuit comprising seven microcircuits (mean of 36.5 million synapses per circuit), with no boundaries between the peripheral circuits and the original microcircuit in the center (only shown in **Figure 2b**). Simulations were run on a BlueGene/Q supercomputer (BlueBrain IV). NEURON models and the connectome are available online at bbp.epfl.ch/nmc-portal^65^.

#### Simulation of in vivo-like spontaneous activity

In the in vivo-like state, release probabilities for all synapses were modulated according to the extracellular calcium concentration found in vivo, leading to substantially lower reliability than in vitro ^16^. As described by Markram *et al.*^17^, the *u_SE_* parameter for synaptic transmission was modulated differentially as a function of extracellular calcium concentration ([*ca*^2+^]_*0*_), allowing transitions from in vitro to in vivo-like dynamics. Neurons were depolarized with a somatic current injection, with currents expressed as a percent of first spike threshold for each neuron, to mimic, for example, the effect of depolarization due to missing neuromodulators. Apart from a small white-noise component (with a variance of 0.001% of the mean injected current per neuron, unless stated otherwise), the current injection was constant. With mean injected currents at around 100% of first spike threshold and [*ca*^2+^]_*0*_ at 1.25 mM, the microcircuit exhibits in vivo-like spontaneous activity ^17^.

#### Simulation of evoked activity

The microcircuit is innervated by 310 (virtual) thalamic fibers^17^. In vivo spike train recordings from 30 VPM neurons were randomly assigned to the 310 fibers, to achieve varying degrees of naturalistic synchronous thalamic inputs. Spike trains were recorded during replayed texture-induced whisker motion in anesthetized rats^39^. Full methods are described in Reimann *et al*.^38^. The second stimulus consisted of synchronous spikes at the 60 central thalamic fibers, with all 60 virtual thalamic neurons firing simultaneously, to approximate a whisker ‘flick’ (see Markram *et al*.^17^).

#### Save-resume

After running a simulation for some amount of biological time, the final states of all variables in the system were written to disk using NEURON’s *SaveState* class. For large-scale simulations, this required the various processes to coordinate how much data each needed to write, so that each rank could then seek the appropriate file offset and together write in parallel without interfering with the others. After restoring a simulation, the user could specify new random seeds (see below).

#### Random numbers

In our simulations, we used random number generators (RNGs) to model all stochastic processes: noisy current injection, stochastic ion channels, probabilistic release of neurotransmitters and generation of spontaneous release events. Each synapse had two RNGs. One was used to determine vesicle release on the arrival of an action potential. The other determined the spontaneous release signal. Similarly, each stochastic *K*^+^-channel model had a RNG determining voltage-dependent opening and closing times. Finally, the white noise process underlying the noisy depolarization was determined by one RNG per neuron. By using different random seeds to initialize the RNGs, we obtained different sequences of random numbers, and consequently different but equally valid simulation outcomes. In earlier versions of the NEURON microcircuit simulation software, the user was given only a single random seed parameter with which to alter the random number streams generated by all RNGs. We added the option to separately change random seeds for RNGs for a specific type of stochastic component. For example, “IonChannelSeed <value>” allows the specification of a seed which is only given to the RNGs used by ion channel instances.

#### Stochastic ion-channels

In some interneuron models, a potassium channel type with a stochastic implementation was added using previously-described methods^17,20,41^. This made it possible to model ion channel noise. Instead of a mean field model, the equations used explicitly track the number of channels in a certain state and allow these numbers to evolve stochastically. When the seed of the random process changes, the small fluctuation caused by the channel noise change with it, but the mean behavior of the ion channel remains the same.

#### Stochastic synapses

The synapse models are described in full detail in Markram *et al*.^17^. Each synapse has one RNG to determine vesicle release upon action potential arrival at the synapse. A second RNG is used to determine the signal for spontaneous miniature post synaptic potentials. When the synapse receives the signal for a spontaneous release event, it is treated as a presynaptic action potential. Therefore, changing the random seed for the minis will eventually change the random number stream for vesicle release for presynaptic spikes, and therefore lead to a “pseudo-deterministic” synapse.

#### Multivesicular release

The synapse model used in this study (see Markram *et al*.^17^) supports multivesicular release (MVR): each release event activates a fraction of the maximal postsynaptic conductance (*g_max_*) proportional to the size of the readily-releasable pool of vesicles (*n_rrp_*). In the univesicular case (*n_rrp_* = 1), the release of one vesicle is sufficient to completely activate the postsynaptic conductance. However, when *n_rrp_* > 1, full activation requires the release of all available presynaptic resources. This allowed us to independently control the mean postsynaptic response to synapse activation (which depends on *g_max_*, but not *n_rrp_*) and its *instantaneous* profile (where *n_rrp_* matters).

#### Deterministic synapse model

In the deterministic synapse model, the *u_SE_* variable is interpreted as the fraction of consumed resources, rather than a release probability. That is, each release event activates a fraction of postsynaptic conductance proportional to *u_SE_*. For this reason, *DetAMPANMDA* and *DetGABAAB* are identical to their stochastic (multivesicular) counterparts in the limit as *n_rrp_* → ∞.

#### Single spike injection

We injected single spikes in twenty different layer 4 pyramidal neurons (and twenty random neurons across the circuit, data not shown) by *replaying* (see below) an additional spike event in one neuron per simulation. Thus, there were no shifted or missing spikes, as may occur when injecting a spike in vivo. The spike was injected *0.1 ms* after resuming the simulation from identical initial conditions.

#### Step-pulse perturbation

We applied a microscopic current step-pulse to all neurons at their soma *0.1 ms* after resuming the simulation (duration: 0.1 ms, amplitude: *1* pA,). The current was chosen to have an almost negligible effect on individual neurons, and was near the limit of the NEURON integrator. On average, 108 ± 8 neurons out of 31,346 neurons had any changes in their spike times (mean of 19 trials ± STD). The majority of the shifted spikes were shifted by less than 0.05 ms (59.1%: < 0.05; 33.1%: < 1 ms.; 5.5%: < 20 ms; 1.8%: < 100 s; 0.5%: < 1 s). Finally, 3 ± 2 neurons had extra or missing spikes. The median first occurrence of an extra or missing spike was at 257 ms (min: 11 ms, max: 946 ms after resuming).

#### Decoupled replay

When resuming a simulation at *t*_0_, we decoupled all connections by setting the connection weights to zero, ensuring that action potentials would be delivered to the synapses of postsynaptic neurons. At the same time, we started *replaying* action potential times from a previous resumed simulation, activating the synapses of postsynaptic neurons as if the presynaptic neuron had fired an action potential, but actually replaying presynaptic action potentials from the previous simulation. For computational reasons, spikes that had not been delivered at the save time *t*_0_ were not delivered in the decoupled replay (meaning that a couple of presynaptic spikes per neuron may have been lost, leading to a slight underestimation of divergence).

### Analysis

#### RMSD and correlation

All analysis was performed using custom scripts written in Python 2.7 using the *NumPy, matplolib* and *SciPy* libraries. Scripts were executed on a Linux cluster connected to the same IBM GPFS file system that the simulation output was written to. Root-mean-square deviation *RMSD_V_* and correlation *r_v_* as defined in Equations 1 and 2 were implemented with *NumPy*.

#### Similarity

The similarity measure *s*(*t*) was defined as the normalized difference between diverging *r_v_* (*t*) (or *RMSD*_v_(*t*)), and steady-state *r_v_*(*t*) (or *RMSD_v_*(*t*)). The steady-state value was defined, as the continuous *r_V,shUffie_*(*t*) computed by shuffling the soma voltages between simulation trials, so that instead of 40 deviating pairs of trajectories, we compared 40 independent pairs of trajectories. As an alternative, we defined it as the mean stationary, fully deviated *r*_∞_ for *t* > *1000 ms* after resuming from identical initial conditions.

#### Firing rate

Firing rate was defined as the average number of spikes in a time interval of size ∆*t*, divided by ∆*t* (∆*t* = 10 ms, unless stated otherwise).

#### Neuron selection

We selected all excitatory neurons in layers 4, 5 and 6 that belonged to the 30 *minicolumns* (out 310 in total) in the center of microcircuit (n = 2024). The analysis was restricted to neurons that spiked at least once in each of the compared simulation paradigms.

#### Spike-timing reliability

Spike-timing reliability was measured using a correlation-based measure first proposed by Schreiber *et al*.^37^. Briefly, the spike times of each neuron in each trial were convolved with a Gaussian kernel of width *σ*_*s*_ = 5 *ms* to yield filtered signals *s*(*n, k; t*) for each neuron n and each trial *k* (∆*t_s_* = 1 ms). The spike-timing reliability for each neuron was then defined as the mean inner product between pairs of signals divided by their magnitude: 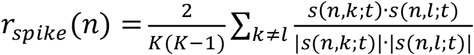, (K = 30; independent trials). Decoupled replay: there are M=5 replays of each of the K=30 trials, and thus 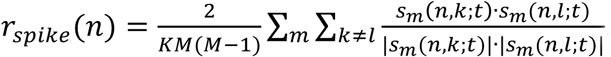.

#### Errors and statistical tests

Error bars and shared areas indicate 95%-confidence intervals (CI), unless stated otherwise. t-based CIs (n = 20; or n = 40 if stated) were computed using *scipy.stats.sem* and *scipy.stats*.*t.ppf* to compute P-values from the CIs. Errors for fit parameters, obtained with *scipy.optimize.curve _fit*, are given as the square-root of the variance of the parameter estimate.

## Author contributions

Conceptualization, M.N., M.R., H.M., E.M.; Methodology, M.N., M.R., H.M., E.M.; Software, M.N., J.K.; Validation, M.N., J.K.; Investigation, M.N; Visualization, M.N.; Writing – Original Draft, M.N., M.R.; Writing – Review & Editing, M.N., M.R., H.M., E.M.; Supervision, H.M., E.M.; Funding Acquisition, H.M.

## Acknowledgements

This work was supported by funding from the ETH Domain for the Blue Brain Project. The Blue Brain Project’s IBM BlueGene/Q system, BlueBrain IV, was funded by the ETH Board and hosted at the Swiss National Supercomputing Center (CSCS). We thank Giuseppe Chindemi, Srikanth Ramaswamy, and Werner Van Geit for help with synapse and ion channel models, and the rest of the Blue Brain team for developing and maintaining the microcircuit model and computational infrastructure. We thank Taylor Newton and Madineh Sedigh-Sarvestani for discussions and critical comments on the manuscript, Richard Walker for extensive feedback on the manuscript, and Oren Amsalem, Mickey London and Idan Segev for helpful discussions.

## Supplementary figures (1 –8)

**Supplementary Figure 1:**
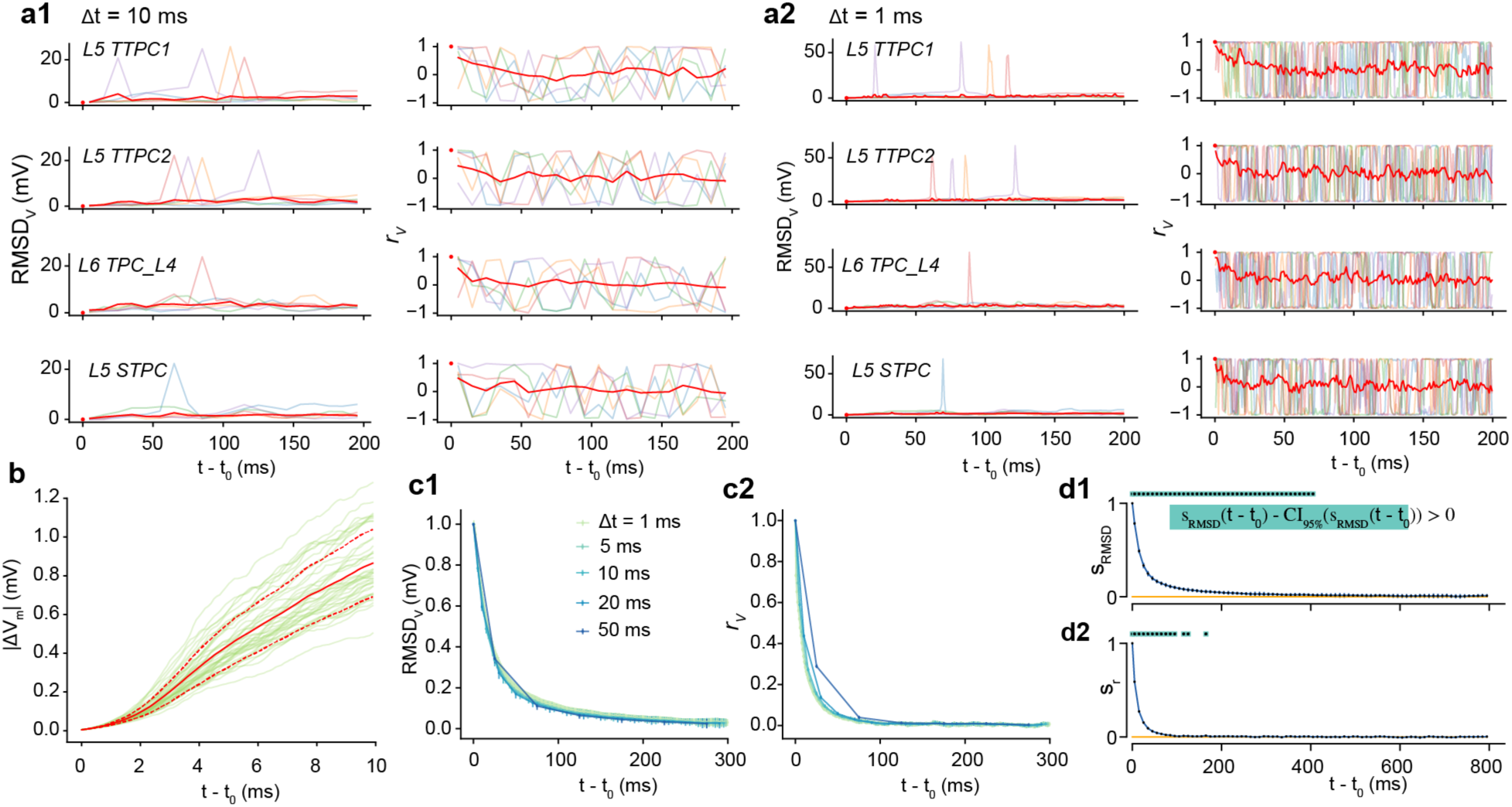
Quantifying the rapid divergence of electrical activity. (**a1**) Root-mean square deviation (*RMSD_V_*) and correlation (*r_v_*) of the somatic membrane potentials between pairs of resumed simulations diverging from identical conditions, for five different base states (faded colors) and the mean of 40 saved base states (red), with ∆*t* = 10 *ms*. Same neurons as in **Fig. 1c**. **a2**) Same as a1, but with ∆*t* = l*ms*. **b**) Mean divergence in the first 10 ms, with ∆*t_v_* = 0.1 *ms* (mean of all neurons and 40 saved base states ± standard deviation). **c**) *RMSD_V_* and *r_v_* for different analysis bin sizes ∆*t*. The time step for the soma voltage is ∆*t_v_* = 0.1 *ms*. **d**) The similarity (*s_RMSD_* and *s_r_*) (mean ± 95% confidence interval). Dots signal where *s_RMSD_* and *s*_r_ are larger than 0, by a 95% confidence interval (p < 0.025).

**Supplementary Figure 2:**
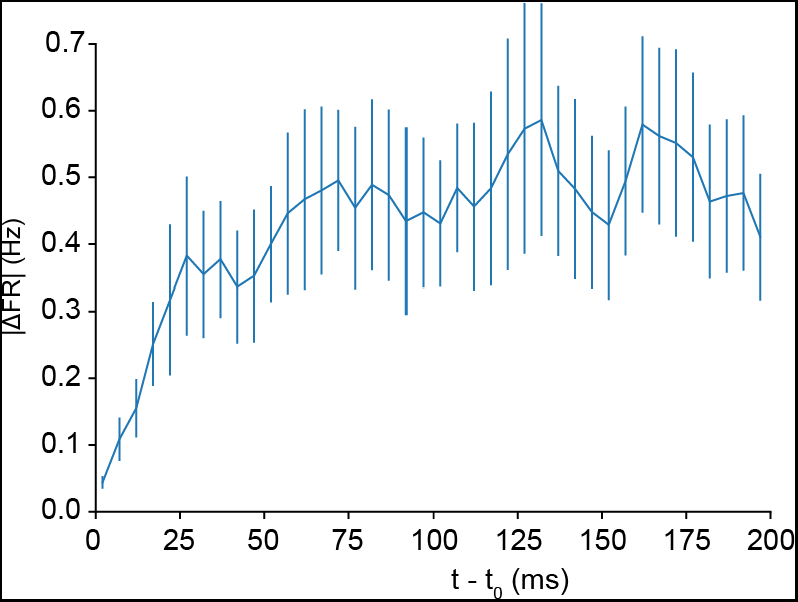
Rapid divergence of population firing rate. Mean population firing rate difference (∆t = 5 ms) between pairs of simulations diverging from identical initial conditions (mean of all neurons and of 40 saved base states ± 95% confidence interval).

**Supplementary Figure 3:**
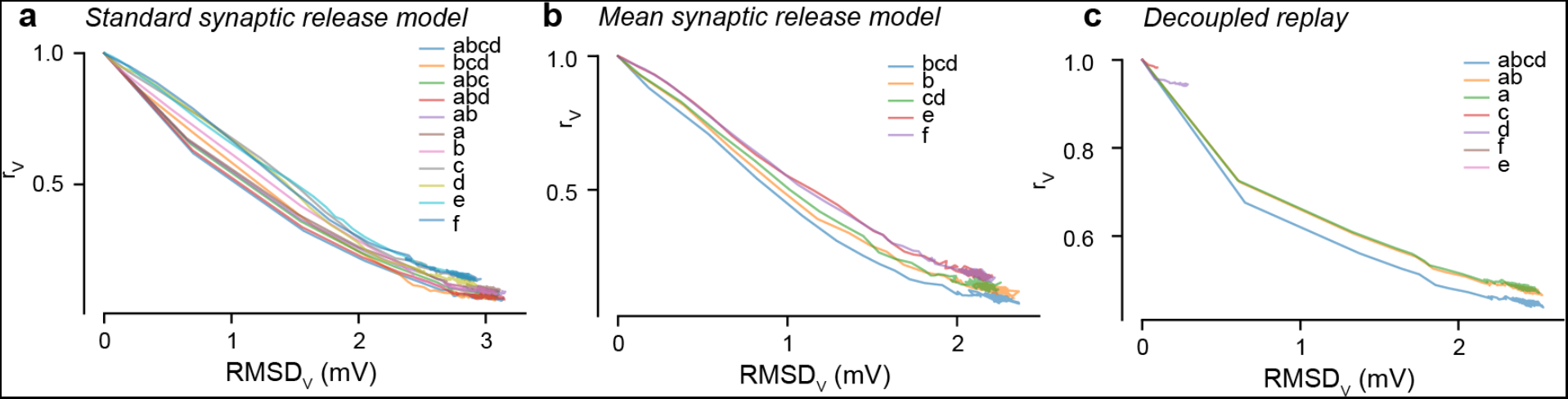
Linear relationship between *RMSD*_v_ and *r_v_*. Mean square deviation (*RMSD*_v,_) and correlation (*r_v_*) of the somatic membrane potentials between pairs of simulations diverging from identical initial conditions (mean of all neurons and saved base states). (**a**) Changing random seeds for subsets of noise sources with the standard stochastic release model. (**b**) Changing random seeds for subsets of noise sources with a mean release model. (**c**) Standard stochastic release model for decoupled, replayed simulations.

**Supplementary Figure 4:**
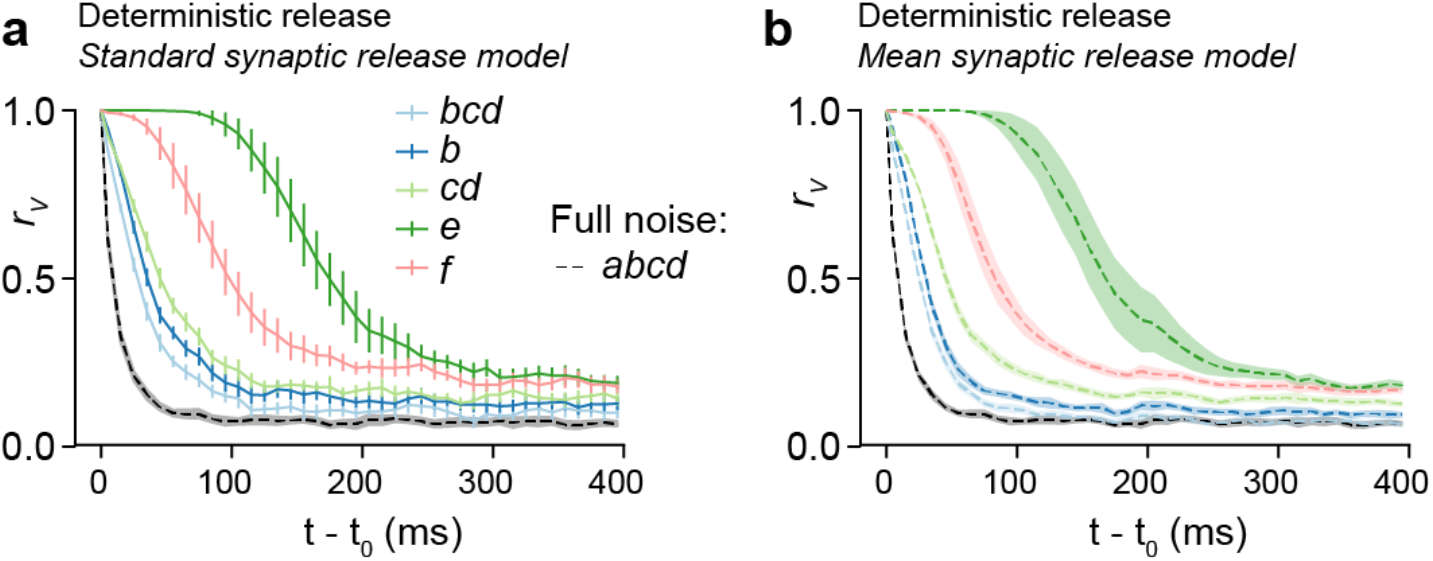
Mean synaptic release model. Correlation *r_v_*, (as in **Fig. 4** and **Supplementary Fig. 5**), with pseudo-deterministic synaptic release by not changing the random seeds for vesicle release (but with a change in ‘mini’ signals for b). (**b**) As in a, but with deterministic synaptic release (mean release model), apart from abcd which has the fully stochastic model.

**Supplementary Figure 5:**
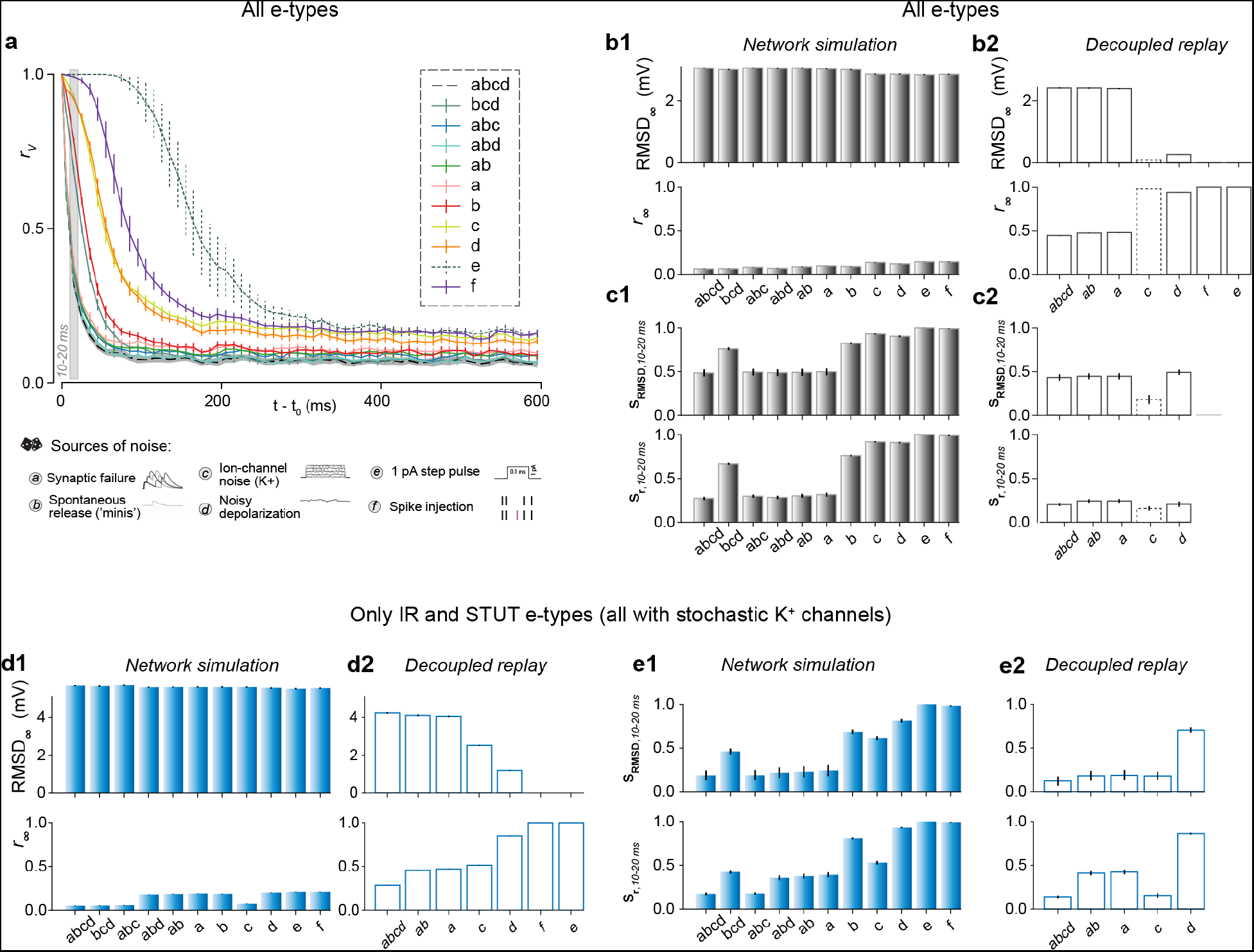
Unravelling noise sources. (**a**) Correlation *r_v_* from identical initial conditions with different cellular noise sources turned on, and when turning of cellular noise, but perturbing the system by a single extra spike (in one neuron) or a miniscule perturbation in all neurons. (**b**) Steady-state membrane potential fluctuations (*RMSD*_∞_) and correlations (*r*_∞_) for network simulations **b1**) and decoupled, replayed simulations **b2**) for different noise sources. **c**) Similarity *s_r_*_/*RMSD*_ at 10-20 ms for network simulations (**c1**) and decoupled, replayed simulations (**c2**) for different noise sources. (**d-e**) Same as **b-c**, but only for the subset of neurons that have stochastic ion-channels (irregularly firing e-types, 1’137 out of 31’346 neurons). All error bars indicate 95% confidence intervals, based on 20 pairs of simulations (40 for *abcd*).

**Supplementary Figure 6:**
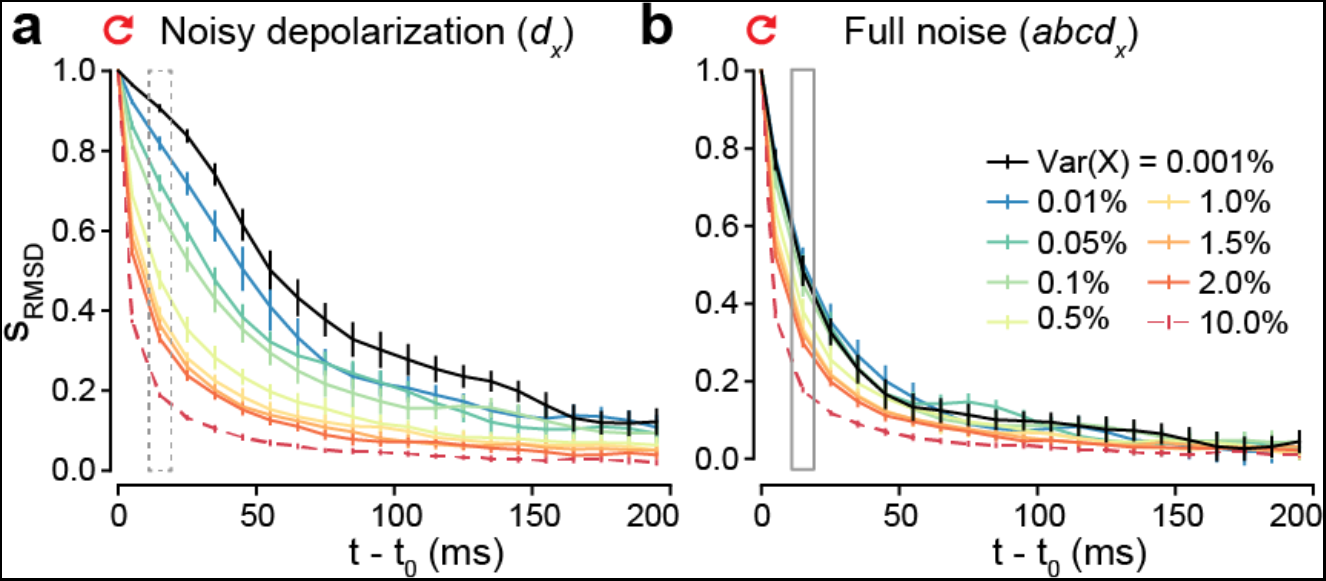
Predicting impact of other noise sources. (**a**) Correlation *r_v_*, when only changing random seeds for noisy depolarization, but with different magnitudes of noise. **b**) As in **a**, but with all noise sources enabled by changing random seeds.

**Supplementary Figure 7:**
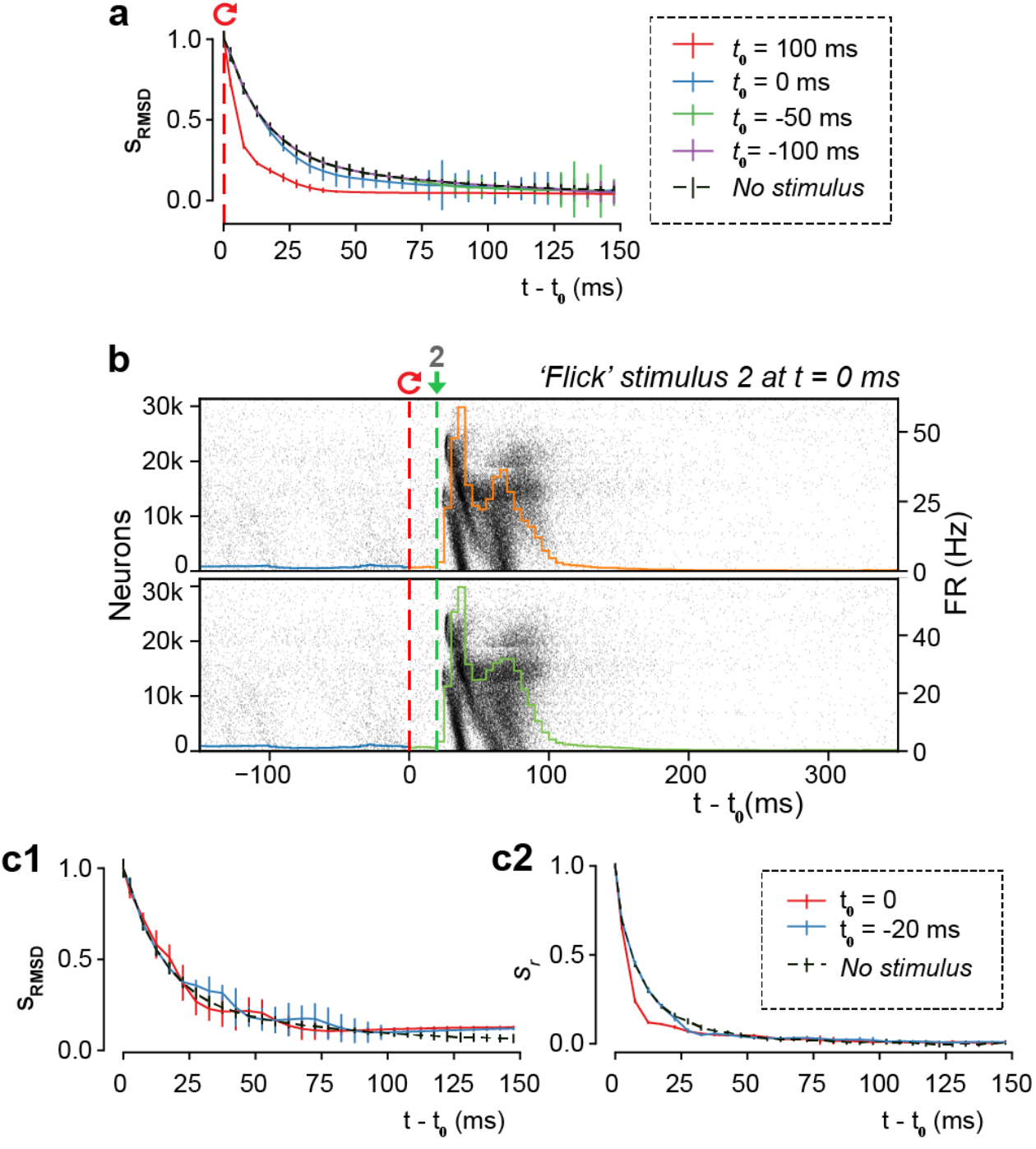
Divergence of evoked activity. (**a**) The similarity *s_RMSD_* defined as the difference between the *RMSD_V_* of diverging and independent trials, normalized to lie between 1 (identical) and 0 (fully diverged) (mean ± 95% confidence interval), for the thalamic stimulus. **b**) Population raster plot and population peristimulus time histogram (PSTH) of all 31’346 neurons in the microcircuit, during evoked activity with a simplified “whisker flick” stimulus (60 VPM neurons are firing at the same time, one spike). (c1) As a, but for the “whisker flick” stimulus. **c2**) As c1, but for *s_r_* instead of *S_RMSD_*.

**Supplementary Figure 8:**
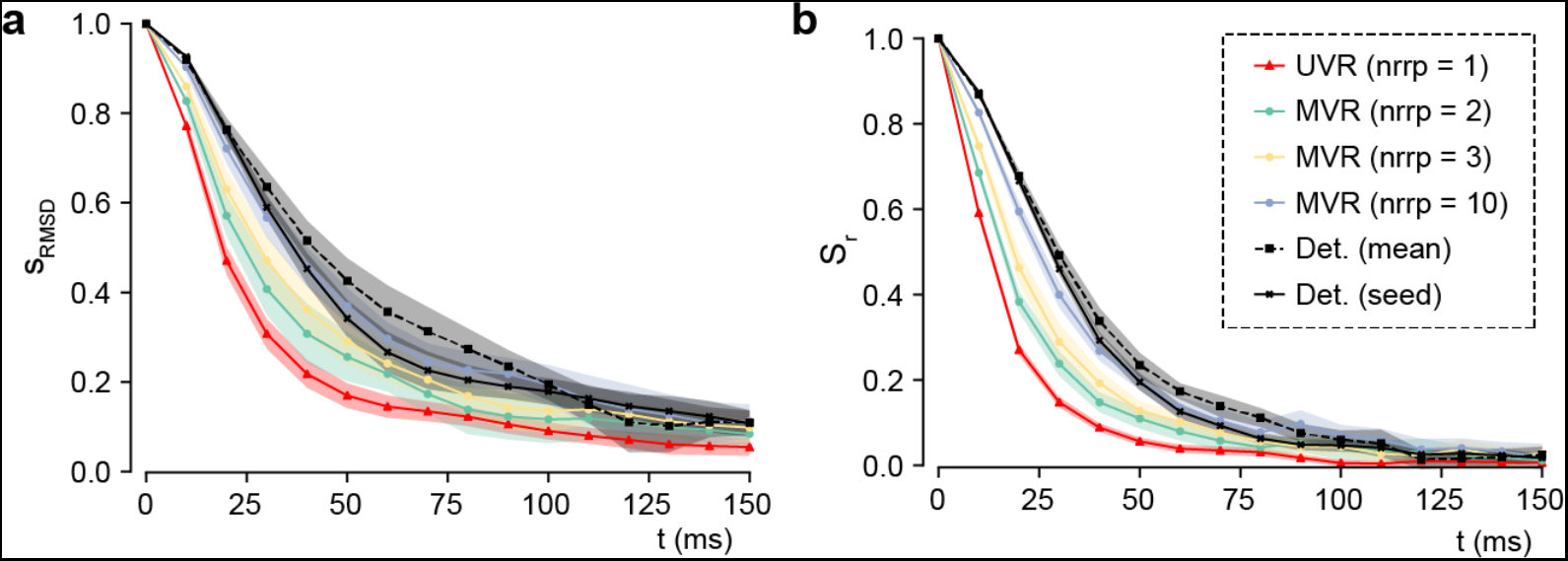
Multivesicular release. Change in divergence time course depending on the size of the pool of readily releasable vesicles (*n_rrp_*). Quantified by similarity of the somatic membrane potentials diverging from identical initial conditions: (**a**) *s_RMSD_* and (**b**) *s_r_*. (mean of all neurons and *n* base states ± 95% confidence interval). (UVR: *n* = 40; all others: *n* = 20).

